# Haemolymphatic tissues of captive boa constrictor *(Boa constrictor):* morphological features in healthy individuals and with Boid Inclusion Body Disease

**DOI:** 10.1101/2024.08.26.609690

**Authors:** E. Dervas, E. Michalopoulou, T. Thiele, F. Baggio, U. Hetzel, A. Kipar

**Affiliations:** Institute of Veterinary Pathology, Vetsuisse Faculty, University of Zurich, Switzerland

**Keywords:** haemolymphatic tissue, Boid Inclusion Body Disease, boa, reptile, involution, granulomatous response

## Abstract

Knowledge on the structure and composition of the haematopoietic tissue (HT) is essential to understand the basic immune functions of the immune system in any species. For reptiles, it is extremely limited, hence we undertook an in-depth in situ investigation of the HT (bone marrow, thymus, spleen, lymphatic tissue of the alimentary tract) in the common boa (*Boa constrictor*). We also assessed age- and disease-related changes, with a special focus on Boid Inclusion Body Disease, a highly relevant reptarenavirus-associated disease in boid snakes. The HT was subjected to gross, histological and ultrastructural examination, including special stains, immunohistochemistry, in situ hybridization and morphometric analyses. In general, the HT was dominated by T cells and lacked a clear structural organization, such as follicle formation. BIBD was associated with significantly higher cellularity and a granulomatous response in the spleen, and the presence of virus-infected haematopoietic cells in the bone marrow, suggesting the latter as a persistent source of viremia.

**Highlights:**

- Lymphatic tissues of *B. constrictor* lack organization and are dominated by T cells
- The thymus of *B. constrictor* shows slow age-related involution
- BIBD-positive *boas* show a granulomatous response in the spleen
- Reptarenaviruses infect haematopoietic cells in the bone marrow
- Reptarenavirus *infected* bone marrow cells might represent a source of viremia

## 1. Introduction

Knowledge on the reptile immune system is extremely limited though it is neither less relevant nor less complex than that of all other animal classes (Zimmerman et al., 2010a). In contrast to mammals, reptiles appear to rely more on the innate immune response than on the adaptive immune system since the latter responds more slowly and is less robust even after repeated pathogen exposure (Zimmerman et al., 2010a). Although this apparent difference in the development of the immune response is not often a focus of research, it is very likely to also have an impact on their response to infectious agents. From an evolutionary perspective, reptiles, as the only living ectothermic amniotes, represent a link between ectothermic anamniotic fishes/amphibians and the endothermic amniotic birds/mammals. Therefore, a better understanding of their immune system may provide important clues regarding the evolutionary history of the immune responses in vertebrates (Vogel et al., 2017).

One of the key elements to decipher the specific features and role of the immune system in any species is to gain insight into the components of the haemolymphatic tissues, their composition and function. In reptiles, these comprise thymus, spleen, lymphatic tissue of the alimentary tract and bone marrow (Cao et al., 2017; den Haan, Joke M M and Kraal, 2012; Hussein et al., 1979; Jacobson and Collins, 1980; Kroese et al., 1985; Kvell et al., 2007; Leceta et al., 1989; Leceta and Zapata, 1985; Oksanen and Tuurala, 1970; Pitchappan and Muthukkaruppan, 1977; Tanaka and Hirahara, 1995). Data on the hematopoietic tissue in snakes is sparse. There is only one morphological study that has tried to provide clues on the mechanism of haematopoiesis in the bone marrow of certain viperid and colubrid snake species (Sano-Martins et al., 2002). Like in mammals, the function, activity and composition of the lymphoid tissue is influenced by various factors like age, sex, stress, diet and reproductive status in reptiles (Andreani et al., 2014; Judson et al., 2020). In reptiles, as ectothermic animals, seasonal temperature variations also have a strong effect (Holding et al., 2014; Jaffredo et al., 2006) which is why several studies have looked into the strong influence of the season on structure and function of spleen and thymus; most describe their atrophy in winter and summer, and activation in spring and autumn. Besides revealing some contradicting data, a literature search highlights substantial knowledge gaps regarding the morphology, function and species-specific differences of the reptilian haematopoietic tissue. For example, in mammals, the thymus is known to involute with age; some authors state a similar involution of the thymus in reptiles whereas others claim its lifelong persistence (Mader, 2009). Several studies mention B cells and immunoglobulins in reptiles (Deza et al., 2007; Gambón-Deza and Espinel, 2008; Zimmerman et al., 2010b), however, the spleen has never been examined morphologically for the presence of B cell specific structures, such as lymphatic follicles (Cohen, 1977; Jaffredo et al., 2006; Vogel et al., 2017). Studies to address such knowledge gaps, and in particular those that would include identification of lymphocyte subpopulations, and especially B cells, in reptiles are hampered by the lack of reliable commercial antibodies that could be applied in *in situ* studies. Due to this limitation the characterization of neoplastic conditions, and in particular round cell tumours, is also often prohibited in reptiles, as e.g. indicated in a study of Schiliger and others, that could not further differentiate a lymphoblastic lymphoma, due to the lack of reliable B cells markers (Schilliger et al., 2011). Also, the assessment of pathological conditions that could affect the immune system, such as infectious diseases, would require the comparative examination of the relevant tissues in healthy and diseased animals and hence knowledge on their normal composition and function.

A prime example for a disease that likely affects the immune system in reptiles is Boid inclusion body disease (BIBD), a widely distributed fatal reptarenavirus-associated disease of boid snakes (Hepojoki et al., 2015; Hetzel et al., 2013; Stenglein et al., 2012). The formation of intracytoplasmic inclusion bodies (IB) comprised of viral proteins, in various tissues and cell types is the morphological hallmark of the disease, however, the presence of IB is not associated with overt cell damage (Baggio et al., 2021; Chang and Jacobson, 2010). BIBD has long been suspected to be associated with immunosuppression, as diseased snakes often develop secondary bacterial, parasitic or fungal infections (Chang and Jacobson, 2010). An older study, undertaken when the causative agent of BIBD was not yet known, even claimed the occurrence of lymphoid depletion in chronically infected animals, unfortunately without substantiating this claim with morphological evidence (Schumacher et al., 1994). In fact, there is indeed no published work that has confirmed changes in the lymphoid system in association with the disease, neither functionally nor morphologically. The present study is a first to address this overt knowledge gap. It describes the normal morphology of the haemolymphatic tissue in *B. constrictor*, and in a second step investigated the same tissues in boas with BIBD for any pathological changes. For this purpose, the haemolymphatic tissue of boa constrictors (*Boa constrictor*) without BIBD and without reptarenavirus infection and with confirmed BIBD were examined by light microscopy including immunohistochemistry and RNA-in situ hybridisation, and by transmission electron microscopy.

## 2. Material and Methods

### 2.1. Study material

The study was undertaken on 59 *B. constrictor* submitted to the Institute of Veterinary Pathology, University of Zurich, for diagnostic post mortem examination. The animals originated from breeding collections and a reptile shelter and were examined upon the owners’ request. In breeding collection A, in Germany, BIBD had previously been diagnosed in several snakes. The owner then had the breeding boas screened for BIBD with the aim to clear his collection from the disease. They then submitted the 26 animals (A1-26) that were diagnosed with BIBD based on the detection of the characteristic IB in blood cells in a peripheral blood smear, an approach applied in previous studies by our group (Alfaro-Alarcón et al., 2022; Argenta et al., 2020; Keller et al., 2017) for euthanasia and post mortem examination, to confirm BIBD and determine any other (associated/secondary/concurrent) pathological conditions. Nineteen animals (B1-B19) originated from a reptile shelter in Switzerland that takes over reptiles from private owners or breeders who are unable to maintain their collection. The snakes were euthanized as the shelter had not succeeded in rehoming the animals within a prolonged period of time. They were subjected to a full post mortem examination to gain information on the general disease status in the shelter (presence of BIBD/reptarenavirus infection or other viral/bacterial diseases of which the owners might not have been aware). For these animals, information on their exact origin was not available anymore. The remaining snakes were from Swiss breeders (C-E) that submitted their animals alive for euthanasia or their euthanised animals, for diagnostic post mortem examination. Information on individual animals and the main post mortem findings are listed in Supplementary Table S1.

### 2.2. Euthanasia, post mortem blood sampling and ethical permission

All snakes submitted alive were euthanized by veterinarians according to the ASPA, Animals (Scientific Procedures) Act 1986, schedule 1 (appropriate methods of humane killing, http://www.legislation.gov.uk/ukpga/1986/14/schedule/1) procedure, and necropsied upon the owner’s request. For anaesthesia 10–20 mg/kg Ketamin (0.1–0.2 ml/kg) and 0.1–0.2 mg/kg Medetomidin (0.1–0.2 ml/kg) were injected intramuscularly in the cranial third of the body. Once deep anaesthesia was achieved, after appr. 45 min, the animal was decapitated. Blood (0.5-3 ml) was collected by cardiocentesis immediately after euthanasia using a 22- or 25-gauge needle on a 3 ml syringe. Two blood smears and four buffy coat samples were prepared based on a previously established protocol (Dervas et al., 2023). For this, the buffy coats were immersed in either 4% formalin for 24 h for light microscopy, or 2.5% glutaraldehyde for 24 h and then buffered in 0.2 M cacodylic acid buffer, pH 7.3 for transmission electron microscopy (TEM). The remaining blood was collected into heparinised tubes (Sarstedt, Germany) and stored at -80 °C until processing.

The study was approved by the institutional review board (MeF-Ethik-2024-01). Also, the terms of service to which owners agree when submitting an animal for a diagnostic post mortem examination include the permission to make use of material from the examination for both teaching and research.

### 2.3. Diagnosis of BIBD

For each animal, one blood smear was stained with May–Grünwald–Giemsa and examined for the presence of IB in blood cells, the standard approach to the intra vitam diagnosis of BIBD that our group has also applied in previous studies (Hetzel et al., 2013; Thiele et al., 2023).

### 2.4. Multiplex polymerase chain reaction for the detection of reptarenavirus infection

All 59 animals in the study were tested for reptarenavirus infection by a multiplex PCR that detects all currently known reptarenaviruses, performed on the full blood and following a recently published method (Baggio et al., 2023). This confirmed the infection in all snakes with BIBD and proved all snakes without BIBD to be free of reptarenavirus infection.

### 2.5. Necropsy and tissue sampling

After euthanasia, snakes underwent a full post mortem examination, including weight and length determination. Samples from all major organs and tissues (brain, respiratory and alimentary tract, liver, kidney, reproductive tract) and from haemolymphatic tissue (thymus, spleen, bone marrow, alimentary tract) were collected and fixed in 10% buffered formalin for appr. 24 h. Additional samples were collected from thymus, spleen and bone marrow and fixed in 2.5 % glutaraldehyde for a minimum of 24 h for transmission electron microscopy (TEM). The bone marrow samples for histology were prepared from formalin or glutaraldehyde fixed vertebrae and ribs. These were lamellated and cross sections prepared using a diamond saw (Exakt 300; Exakt) and, in the case of adult snakes, followed by gentle decalcification in RDF (Biosystems) for 1-2 months at room temperature (RT) and on a shaker. In the case of juvenile snakes (<6 months of age), decalcification was not necessary, and cross sections of ribs and vertebrae were directly embedded.

Since none of the animals exhibited overt gross changes, further examinations (bacteriological or parasitological examination) were not performed. The extent and distribution of connective tissue in the haematopoietic tissue (thymus, spleen, bone marrow) was determined based on different special stains. The Van Giesson stain served to highlight collagen fibres. After deparaffinisation and rehydration, sections were incubated with the Weigert’s Haematoxylin Kit (Waldeck GmbH & Co, Münster, Germany) according to the manufacturer’s protocol, followed by staining in picrofuchsin solution (Waldeck GmbH & Co) for 5 min. For staining of elastic fibres, sections were stained with a combination of resorcinol and fuchsine (Waldeck GmbH & Co) for 20 min, then counterstained for 5 min using Nuclear fast red-aluminium sulfate solution (0.1%) (Sigma-Aldrich, Buchs, Germany). For the identification of reticulin fibres, the Reticulin/Nuclear Fast Stain Kit (Agilent Dako, Germany) was applied on the Artisan Link and Artisan Link Pro Staining Systems (Agilent Dako). For the detection of ferric ion in tissues, the deparaffinized and rehydrated sections were stained in 2% Hydrochloric Acid (Huber Lab, Aesch, Switzerland) and 2% Potassium Ferrocyanide (Sigma-Aldrich) (1:1). Afterwards, they were counterstained for 5 min with Nuclear fast red-aluminium sulfate solution (0.1%). For the detection of fibrin, the deparaffinized and rehydrated sections were stained with chromogen solution (potassium dichromate 3%, hydrochloric acid 10%) for 30 min, followed by staining with a differentiating solution (potassium permanganate 0.5%, sulfuric acid 3%) for 1 min and bleaching with oxalic acid (1 %) for 1 min. After rinsing with tap water, the slides were left in the Phosphotungstic acid-haematoxylin stain (PTAH) solution (haematoxylin 0.1% and phosphotungstic acid 2%) overnight. As a final step, the sections were dehydrated, cleared in xylene, and coverslipped (HistoCore Spectra CV, Leica, Buffalo Grove, USA).

### 2.6. Immunohistochemical staining and RNA in situ hybridization (RNA-ISH)

Commercial antibodies designed to target protein epitopes of leukocytes or structural components that can be used in immunohistological examinations are very limited in snakes, and reptiles in general. Therefore, to establish suitable staining protocols for use in B. constrictor, a range of antibodies and protocols established for other reptile species or other animal classes were tested on sections from the haemolymphatic tissue of some snakes. For antibodies that showed cross-reactivity and in which the protein and/or mRNA sequences of the antigens were known, we tested their percentage of identity using BLAST analysis (NCBI, BLAST) to the *Python bivitattus* genome (accession number XM_015890515.2).

These test runs resulted in the immunohistochemical staining protocols for the following cells and proteins: T cells (CD3+), monocytes/macrophages (Iba1+), vascular endothelial cells (factor VIII–related antigen; Factor VIII+), epithelial cells (Pan-Cytokeratin 26 (PCK26)), smooth muscle cells (α–smooth muscle actin; α-SMA+) and apoptotic cells (cleaved caspase-3+). These were applied to sections from haemolymphatic tissues of selected animals (Supplemental Table 1). For the detection of reptarenaviral nucleoprotein (NP) as well as IgM and IgY expressing cells, custom-made rabbit polyclonal antibodies were used, following previously published protocols (Hetzel et al., 2013; Korzyukov et al., 2016). For all immunohistochemical protocols, the horseradish peroxidase method was applied, using a Dako autostainer (Agilent Dako).

Primary and secondary antibodies and the respective antigen retrieval and detection methods are listed in Table 1. Briefly, after deparaffinization, antigen retrieval was performed for all antigens except for most antigens, by incubation of the slides in citrate buffer (pH 6) or EDTA buffer (pH 9) at 98 °C for 20 min. After incubation with the primary antibodies, endogenous peroxidase was blocked by incubation with hydrogen peroxide solution for 10 min at RT. Afterwards, sections were incubated with the secondary antibodies, according to the manufacturer’s protocol. Sections were washed with phosphate buffered saline (PBD; pH 7.4) between each incubation step. Finally, sections were counterstained with haematoxylin for 40 s and mounted. Sections incubated with non-immune serum from the species in which the primary antibody was raised instead of the primary antibodies served as negative controls.

**Table 1.**
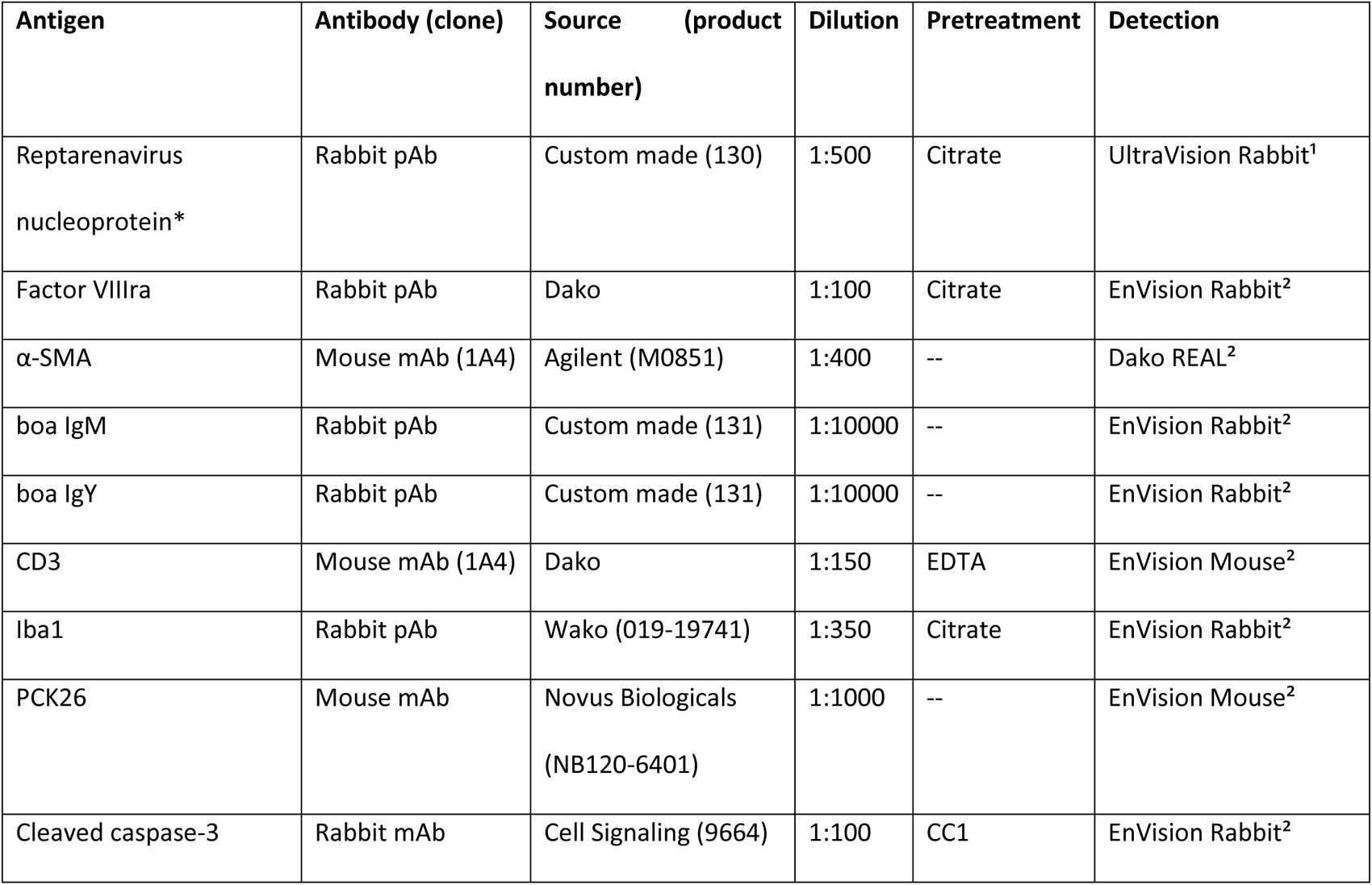
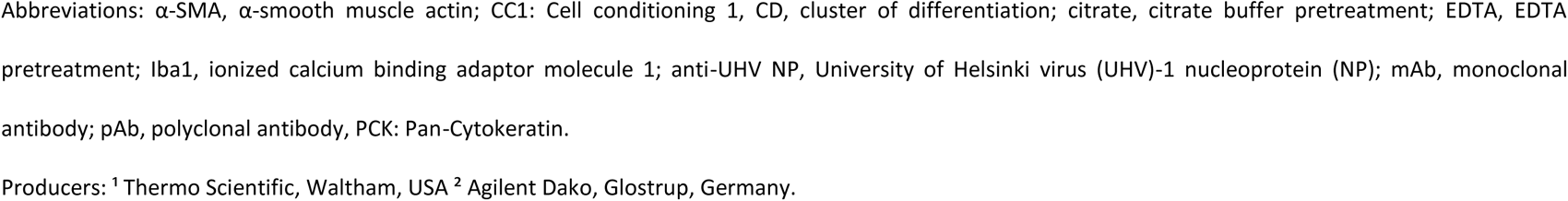
Antibodies, antigen retrieval and detection methods applied for immunohistochemical staining.

| Antigen | Antibody (clone) | Source (product number) | Dilution | Pretreatment | Detection |
| --- | --- | --- | --- | --- | --- |
| Reptarenavirus nucleoprotein* | Rabbit pAb | Custom made (130) | 1:500 | Citrate | UltraVision Rabbit<sup>1</sup> |
| Factor VIIIra | Rabbit pAb | Dako | 1:100 | Citrate | EnVision Rabbit<sup>2</sup> |
| α-SMA | Mouse mAb (1A4) | Agilent (M0851) | 1:400 | -- | Dako REAL<sup>2</sup> |
| boa IgM | Rabbit pAb | Custom made (131) | 1:10000 | -- | EnVision Rabbit<sup>2</sup> |
| boa IgY | Rabbit pAb | Custom made (131) | 1:10000 | -- | EnVision Rabbit<sup>2</sup> |
| CD3 | Mouse mAb (1A4) | Dako | 1:150 | EDTA | EnVision Mouse<sup>2</sup> |
| Iba1 | Rabbit pAb | Wako (019-19741) | 1:350 | Citrate | EnVision Rabbit<sup>2</sup> |
| PCK26 | Mouse mAb | Novus Biologicals (NB120-6401) | 1:1000 | -- | EnVision Mouse<sup>2</sup> |
| Cleaved caspase-3 | Rabbit mAb | Cell Signaling (9664) | 1:100 | CC1 | EnVision Rabbit<sup>2</sup> |

Abbreviations: α-SMA, $\alpha$ -smooth muscle actin; CC1: Cell conditioning 1, CD, cluster of differentiation; citrate, citrate buffer pretreatment; EDTA, EDTA pretreatment; Iba1, ionized calcium binding adaptor molecule 1; anti-UHV NP, University of Helsinki virus (UHV)-1 nucleoprotein (NP); mAb, monoclonal antibody; pAb, polyclonal antibody, PCK: Pan-Cytokeratin.
Producers: <sup>1</sup> Thermo Scientific, Waltham, USA <sup>2</sup> Agilent Dako, Glostrup, Germany.

Unfortunately, a range of commercial antibodies against human and/or murine proteins or peptides for the identification of B cells (e.g. CD20, CD79a, PAX-5) did not yield any reaction in snake tissues when various immunohistochemistry protocols. We therefore used an RNA-ISH approach, and specifically the RNAscope® technology (Advanced Cell Diagnostics Inc (ACD), Newark, USA), and applied oligoprobes coding for *B. constrictor* CD20 in order to detect B cells in the snake tissue, following previously established protocols (Dervas et al., 2023).

### 2.7. Morphometric analyses

A morphometric approach was taken to quantify the overall cellularity as well as the number/percentage of macrophages (Iba1+), T cells (CD3+) and apoptotic cells (CC3+) in spleen and thymus. Slides stained with HE, for Iba1, CD3 and CC3 were scanned using a digital slide scanner (NanoZoomer-XR C12000; Hamamatsu, Hamamatsu City, Japan) and evaluated with the computer programme VIS (Visiopharm Integrator System, Version 5.0.4. 1382; Visiopharm, Hoersholm, Denmark). For all quantitative approaches, entire cross sections of thymus and spleen were manually selected as ROI. In cases where more than one cross section of an organ was present on the slide, the quantitative assessment was performed on all sections and the means of the measurements of each organ were calculated and used for statistical analysis. Sections stained with HE served to assess the total number of nuclei and thereby the overall cellularity of spleen and thymus. A decision forest classification method was applied and a number of post-processing steps (change by shape (cell diameter <4 μm^2^, object separation (surrounding width of 1 μm) allowed recognition and separation of individual cells. Bigger vessels were excluded manually from the ROI to avoid counting of erythrocytes as these are nucleated cells in snakes and could therefore be counted as parenchymal cells. Results were presented as the total nuclei count per ROI area (in μm2).

Sections stained for Iba1 were used to quantify the total Iba1-positive area (corresponding to monocytes/macrophages) in spleen and thymus. A threshold classification method allowed recognition of the Iba1-positive area, and the results were expressed as the percentage of positive area (in μm^2^) per total area of the ROI (in μm^2^). In a post-processing step, very small positive areas (<5 μm^2^) were excluded from counting to avoid falsely classifying areas of increased background staining as cells. Sections stained for CD3 were used to quantify the total CD3-positive area (T cells) in spleen and thymus. The CC3 stain served to determine the number of cells undergoing apoptosis. A cell classification method combined with a manually tuned classification of the detection of a cytoplasmic immunoreactivity allowed recognition of positive (intense brown staining of the entire cytoplasm) and negative cells in each ROI, and the results were expressed as the absolute number of positive cells per ROI area. In a post-processing step, an intensity threshold of <120 was applied for all markers to avoid falsely classifying heterophils that routinely show an intensely green staining of their granules after any immunohistological staining (Dervas et al., 2023).

### 2.8. Transmission electron microscopy (TEM)

For TEM, tissue samples (spleen, thymus, bone marrow) from selected animals and a few buffy coat samples from selected animals (Supplemental Table 1) were fixed for 24 h in 5% glutaraldehyde, buffered in 0.2 M cacodylic acid buffer, pH 7.3, then trimmed and routinely embedded in epoxy resin. Semithin sections (1.5 μm) were prepared from the epoxy resin blocks and stained with Toluidin blue. These served to select areas of interest for the preparation of ultrathin sections (75 nm) that were contrasted with lead citrate and uranyl acetate and viewed with a Philips CM10, operating with a Gatan Orius Sc1000 digital camera (Gatan Microscopical Suite, Digital Micrograph, Pleasanton, USA).

### 2.9. Statistical analysis

All morphometric parameters were analysed using Stata 13 (StataCorp. 2013. Stata Statistical Software: Release 13. College Station, TX: StataCorp LP.). The level of significance testing was set with a P value of 0.05. Descriptive statistics were applied, and the data were tested for normality by the D’Agostino-Pearson normality test. Since all data was normally distributed, an independent sample t-test was used to determine differences between BIBD-positive and BIBD-negative animals. Correlations of the overall cell density of the thymus and spleen with the age of each animal were evaluated using the Spearman’s rank correlation coefficient. The size of the correlation coefficient was interpreted as “low” for correlation coefficients lower than 0.5, “moderate” for correlation coefficients between 0.5 and 0.7, and “high” for correlation coefficients higher than 0.7 (Mukaka, 2012). Geometric and arithmetic mean, including confidence intervals, were determined for logarithmically transformed and raw data respectively. Variables where data were normally distributes include the splenic and thymic apoptotic cells, total splenic and thymic cell number, thymic T cells and macrophages. Logarithmic transformation was applied on splenic macrophages and T cells.

### 2.10. Morphological identification of structural components and cell types in haemolymphatic tissues and age group allocation

The haematopoietic tissue of animals without evidence of BIBD (see 2.3) and confirmed negative for reptarenavirus infection (see 2.4), hereafter called “control animals”, served for morphological identification of structural components and cell types.

In general, studies on the morphological components of haematopoietic tissue and peripheral blood cells of reptiles, and *B. constrictor* in particular, are sparse, often limited to a single species, and to a certain degree inconsistent in their terminology. Therefore, to avoid confusion, provide some consistency and allow future comparative studies, we decided to use the nomenclature applied for anatomical structures/locations in the relevant literature so far, to provide as much consistency and transparency as possible, for future studies and comparative use.

Bone marrow: For vertebrae and medullary cavities, the anatomic terminology introduced in two publications for the bone marrow of various snake species, including *B. constrictor*, was employed ((Sano-Martins et al., 2002) (Raül Carmona et al., 2010). The ultrastructural identification of bone marrow cells was based on our recent work on peripheral blood cells of *B. constrictor* (Dervas et al., 2023), older studies on the ultrastructure of the bone marrow in snakes, lizards and birds (Campbell and Grant, 2021; Dabrowski et al., 2007; Zapata et al., 1981) , and reports on peripheral blood cells of other reptile species (Moller et al., 2016; Salakij et al., 2002). In general, precursor cell types of all hematopoietic stem cell lines were recognized due to their morphological similarities to their mature counterpart (e.g. size, cytoplasmic processes, cell organelles like granules) and their location (e.g. erythropoietic precursors are mainly found in close proximity to the sinus venosus (Dabrowski et al., 2007)). In humans and many mammals, the function and morphology of cells in all stages of haematopoiesis has been described in great detail (e.g. erythropoiesis is divided into 5 stages: proerythroblasts, erythroblasts, normoblasts, reticulocytes and mature erythrocytes) (Deldar et al., 1989; Kashimura, 1982; Pease, 1956). Since we could not identify similar stages with certainty in the bone marrow of *B. constrictor*, and to avoid overinterpretation, we applied the prefix “pro” to address the respective cells in the bone marrow (proerythrocytes, promonocytes etc.). Since the approach of the present study does not allow further comments on the functional or morphological level of maturation of the bone marrow cells, the terms “immature” or “mature” were avoided.

Due to the structural similarity of the spleen, we applied the terminology, and in particular the term “perilymphoid fibrous zone” (PLFZ) introduced for the spleen of an elaphid snake (Yasukazu Tanaka and Yoshihiro Hirahara, 1995) to the spleen of *B. constrictor*, complemented by the terminology introduced for structural components of spleen of other reptile species; this also applies to the thymus (Hussein et al., 1979; Leceta and Zapata, 1985; R. E. Ridi et al., 1981; Wang et al., 2022; Yasukazu Tanaka and Yoshihiro Hirahara, 1995). For the lymphatic tissue in the alimentary tract, morphological studies in different reptile species, e.g. including Argentine tegus, elaphid and boid snakes, were considered (Betancourt et al., 2022; Jacobson and Collins, 1980; R. E. Ridi et al., 1981).

To allow determination of age-related differences in the haemolymphatic organs, two age classes were established, following an approach we have previously taken (Dervas et al., 2023): animals < 3 years of age (prior to definite sexual maturity and reproduction; “young”), and animals ≥3 years of age (sexually mature, reproducing animals; “adult”).

## 3. Results

### 3.1. Study population and pathological changes

Among the 59 animals included into the study, 26 did not exhibit IBs in blood cells and were negative for reptarenavirus infection (“control animals“). Of these, 8 were female and 10 male; for 8 juvenile animals the sex could not be determined due to lack of differentiation of the gonads. The age was known for 25 snakes; it ranged from 3 months to 10 years, with a mean of 3.58 years; 15 were “young” (< 3 years of age) and 11 “adult” (≥3 years of age). None of the animals had presented clinical signs prior to euthanasia. Pathological changes were rare and restricted to a moderate chronic oesophagitis (animal B10), a moderate lymphocytic (animal A23) and mild granulomatous pneumonia (animal B7).

Thirty-three animals were BIBD-positive, with confirmed reptarenavirus infection; 20 were female, and 13 male. The age was known for 32 snakes; it ranged from 1 year to 25 years, with a mean of 4.59 years; 16 were categorized as “young” and 17 as “adult”. The majority of animals had not presented clinical signs prior to euthanasia. Two had shown weight loss (A10, A13), two diarrhoea (A15, A16), and one respiratory distress (B16). Overall, histological changes were observed in 27 of the 33 BIBD-positive necropsied animals (81%; excluding changes in haemolymphatic tissues). The most common finding was heterophilic (entero-)colitis of variable severity (23/33 animals; 69%). Individual animal data are provided in Supplemental Table S1.

### 3.2 Bone marrow

#### 3.2.1 Location, morphological features and composition

Haematopoietic tissue was found within variably sized medullary cavities (lacunae) of the cervical, trunk and cloacal vertebrae, and in the ribs (distally of the costal head) regardless of the animals’ age. In the vertebrae, lacunae were found in the following locations: 1. the dorsal part of vertebrae in the so-called neural spine; 2. bilaterally in the neural arch in the postzygapophyseal and transverse processes and the parapophysis; 3. in the central bony part and peripheral cartilaginous part of the vertebral centre (Fig. 1A and B).

**Fig. 1.**
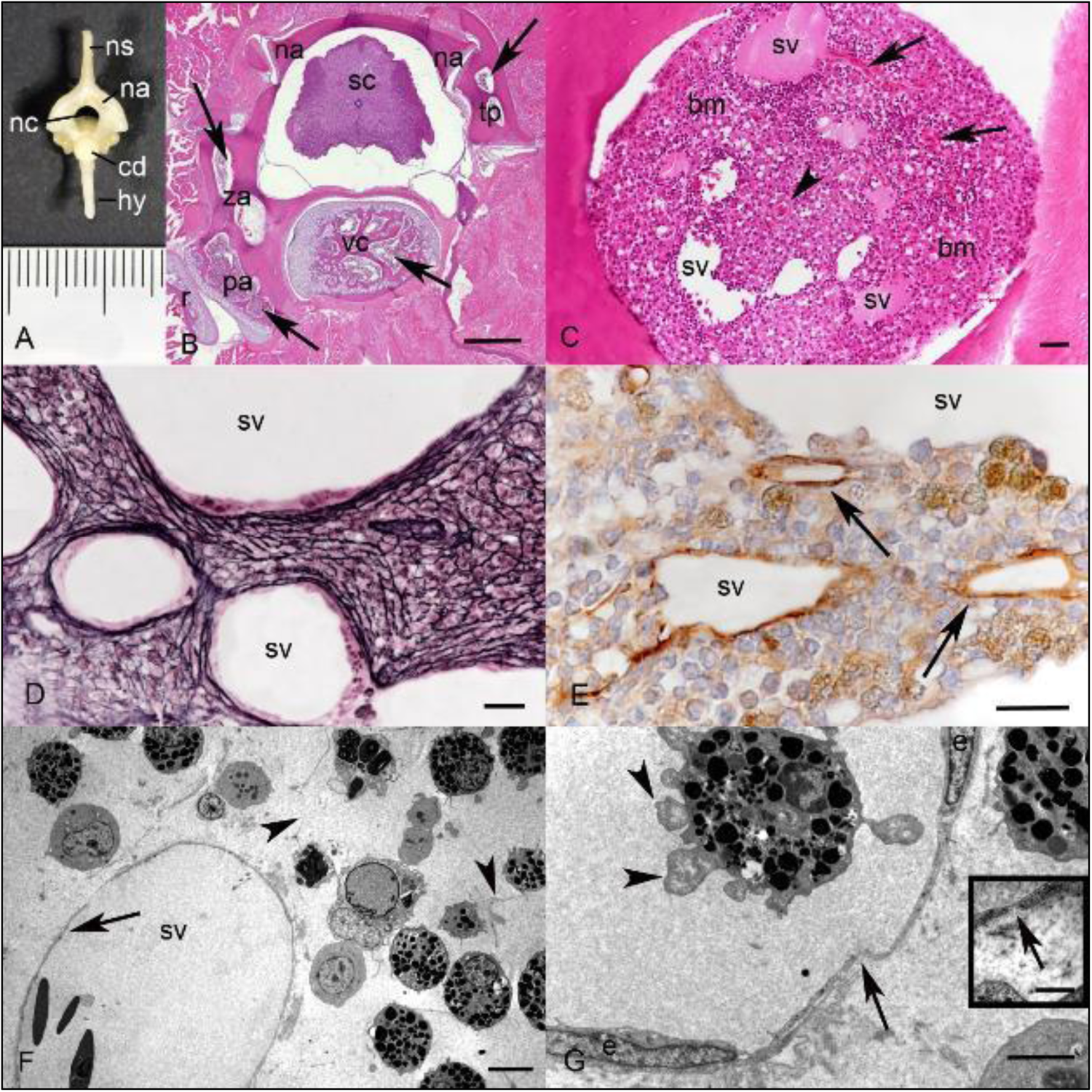
Anatomic features of the bone marrow in *B. constrictor*. **A.** Cervical vertebrae of an adult *B. constrictor* (animal C3, 1 year). ns: neural spine; na: neural arch; nc: neural canal; cd: condyle; hy: hypapophysis. **B. Vertebral cross section from the trunk part** (animal B1, 8 years). On both sides of the neural arch (na), the transverse processes (tp) and the postzygapophysial processes (za) are filled with bone marrow (arrows). In the vertebral centre (vc) and the parapophysis (pa) numerous lacunae are filled with bone marrow (arrow). Lacunae with bone marrow are also found in the rib (r), starting from the costal head. sc: spinal cord. HE stain, bar = 250 µm. **C. Costal bone marrow (bm)** (animal A4, 2 years). Multiple sinus venosus (sv) containing proteinaceous fluid and cells (erythrocytes, haematopoetic cells), central artery (arrowhead) and multiple small venules (arrows). HE stain, bar = 20 µm. **D. Connective tissue** (animal A19, 11 years). A dense meshwork of reticulin fibres extending from the sinus venosus (sv) supports the haematopoietic cells. Reticulin stain, bar= 20 µm. **E. Vascular structures** (animal A3, 8 years). Staining for the endothelial cell marker Factor VIII highlights the presence of multiple variably sized sinus venosus (sv) and small venules (arrows), covered by a thin layer of endothelial cells. Immunohistochemistry, haematoxylin counterstain, bar = 20 µm. **F. Haematopoietic cells adjacent to a sinus venosus in the rib of a *B. constrictor*** (animal B17, 3 months). The sinus venosus (sv, arrow) contains a few erythrocytes and is surrounded by a thin meshwork of reticulin fibres (arrowheads) with embedded haematopoietic cells. TEM, bar = 5 µm. **G**. **Wall of the sinus venosus** (animal B17, 3 months). The sinus venosus (sv) is lined by a layer of flat endothelial cells (e). Inset (higher magnification of area highlighted by arrow): Endothelial cells are supported by a discontinuous basement membrane (arrowhead) that leads to the formation of small openings (pores, arrow). The granulocyte in the lumen of the sinus venosus has cytoplasmic projections (pseudopodia, arrowheads) that enable locomotion. TEM, bar = 2 µm.

The main vascular structures in the marrow cavity were multiple variably sized venous sinuses (*Sinus venosus*); these were filled with erythrocytes and/or mature peripheral blood cells, like granulocytes and monocytes. A few small-caliber arterioles, distinguishable by their tunica media, and small venules, lacking a tunica media, were also noted (Fig. 1C). The vascular structures were embedded in a dense meshwork of fine reticular fibres, supported by few elongated spindeloid reticular cells (Fig. 1D). The venous sinuses were lined by a thin monolayer of endothelial cells (Fig. 1E and F). The ultrastructural examination showed that the endothelial cells, characterised by elongated, central nuclei and scant cytoplasmic organelles (mitochondria, occasional lysosomes), were supported by a discontinuous basement membrane with pores (fenestration, Fig. 1G).

The overall cellularity and cellular composition of the haematopoietic tissue showed mild, yet noticeable age-associated differences. In young animals, a lower cellularity and a predominance of granulocytic precursor cells was observed (Fig. 2A); this was particularly evident in the youngest animals (3 months of age). In adult individuals, the cellularity was higher, as the entire marrow cavity was generally densely packed with cells (Fig. 2B). There was also a higher number of cells resembling erythropoietic cells or cells with azurophil and monocyte morphology (both the latter being Iba1 positive, as shown in a recent publication (Dervas et al., 2023)). It should be noted that, as a consequence of the prolonged exposure to the formic acid containing decalcifying solution necessary to decalcify the bones, cellular structures were difficult to discern due to the reduced differentiation in HE stained sections.

**Fig. 2.**
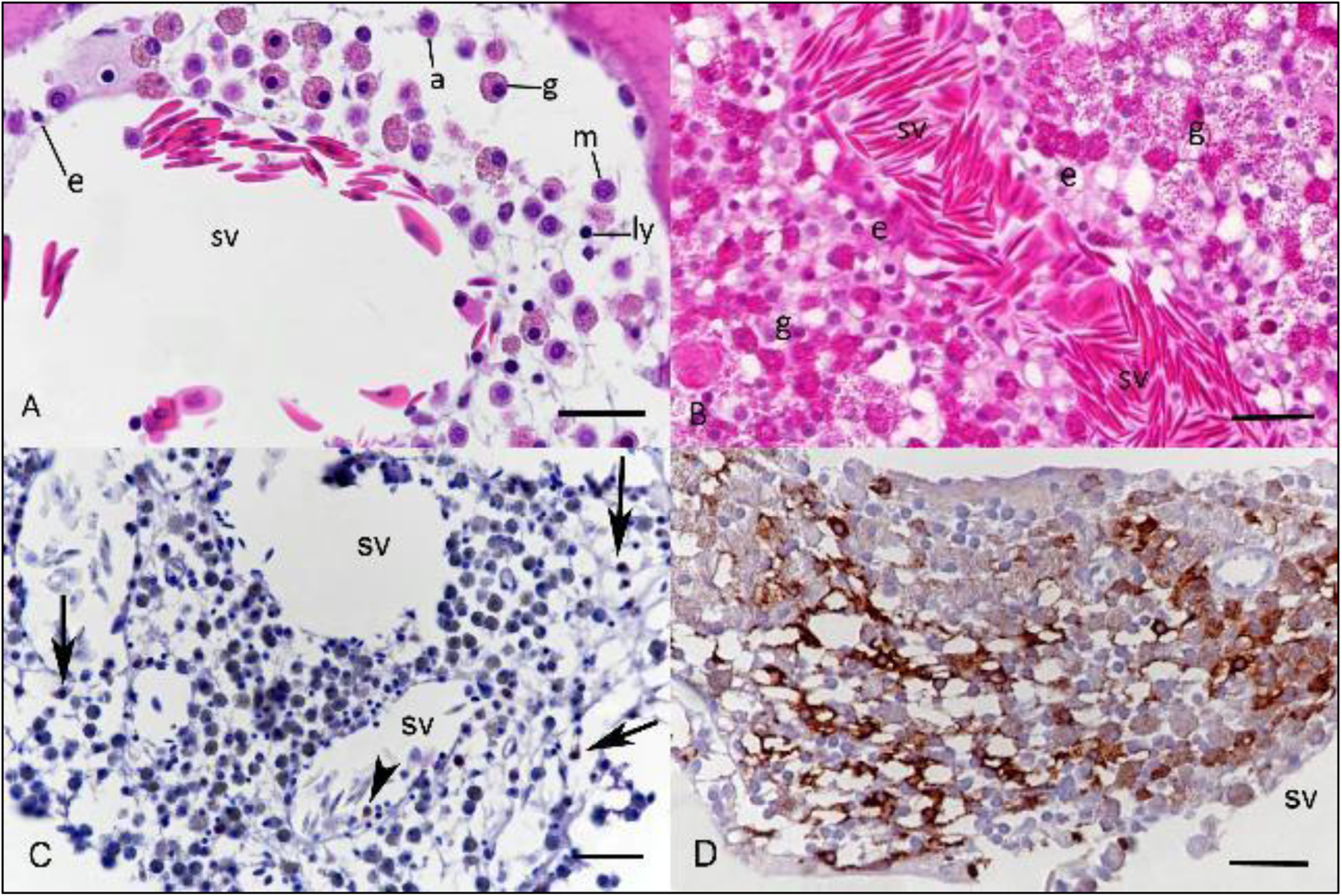
Composition of the bone marrow in *B. constrictor*. **A.** Vertebral bone marrow of a young animal (animal B17, 3 months), exhibiting low to moderate cellularity. The sinus venosus (sv) is filled with erythrocytes and rare mature haematopoietic cells. The extravascular space around the sinus venosus contains a few small proerythrocytes (e), several progranulocytes (g) with fine cytoplasmic granules, promonocytes (m), proazurophils (a) and a few prolymphocytes (ly). Proazurophils can be distinguished from promonoytes by their brighter eosinophilic cytoplasm. HE stain. **B. Costal bone marrow of an adult animal** (animal B1, 8 years), exhibiting high cellularity. Proerythrocytes (e) are located around the sinus venosus (sv), whereas progranulocytes (g) are mainly present in its periphery. HE stain. **C, D. Iba1 positive cells in the bone marrow of young animals. C. Costal bone marrow** (animal B17, 3 months) with rare positive cells (arrows). **D. Vertebral bone marrow** (animal A11, 2 years), with numerous positive cells. Iba1 positive cells are interpreted as promonocytes and proazurophils (Dervas et al., 2023). Immunohistochemistry, haematoxylin counterstain. Bars = 20 µm.

<u>Erythropoiesis</u>. Erythropoietic precursor cells (proerythrocytes) were found outside the sinus venosus, densely packed around its wall. In HE stained sections, the cells were small and round, with a hyperchromatic dark round central nucleus and a thin rim of homogenous, strongly eosinophilic to dark basophilic cytoplasm (Fig. 2A and B). Ultrastructurally, proerythrocytes were characterised by a round central nucleus with a moderate amount of clumped heterochromatin and a ribosome-rich cytoplasm with few scattered organelles (mitochondria, endoplasmic reticulum (ER) (Supplemental Fig. 1A).

<u>Granulopoiesis</u>. Most of the remaining haematopoietic space, surrounding the venous sinuses, was tightly filled with granulopoietic cells, comprising larger round cells with abundant cytoplasm that was densely packed with multiple variably eosinophilic, often birefringent granules, and a round central or eccentric hyperchromatic nucleus (Fig. 2A and B). Ultrastructurally, progranulocytes were round cells with a regular outline that were mainly located in the centre of the medullary cavity. Their cytoplasm was packed with granules of variable size and electron density. Granules of 0.3 - 0.5 μm size and low electron density, interpreted as primary granules, and larger (1-1.5 μm) round or polygonal granules with electron-dense cores, interpreted as secondary granules, were both abundant; there were also lesser medium sized (0.8 -1 μm) granules of high electron density, interpreted as tertiary granules. A moderate number of cytoplasmic organelles, such as mitochondria, were also observed (Supplemental Fig. 1B). The nucleus showed clumped, often peripherally orientated heterochromatin. (Pro)granulocytes associated with the sinus venosus, both in the lumen and in the periphery, with a high number of both small cytoplasm-filled membrane projections (these are known to mediate locomotion (Van Haastert, Peter J M and Devreotes, 2004)) and cytoplasmic granules, in the majority consistent with tertiary granules, were interpreted as more mature granulocytes (Sano-Martins et al., 2002). These cells also displayed a noticeably lesser amount of cell organelles, and in particular only few mitochondria (Supplemental Fig. 1C). Granulocyte subtypes, such as heterophils, basophils and eosinophils, could not be distinguished in the bone marrow with certainty. Whilst they contained primary, secondary and tertiary granules, they did not exhibit distinct features that would allow designation to a granulocyte subtype.

<u>Monocytopoeisis, monocytes and azurophils</u>. In HE-stained sections, cells with monocyte and azurophil morphology were readily distinguishable from progranulocytes due to their smaller size and the absence of prominent intracytoplasmic granules (Fig. 2A) (Dervas et al., 2023). Immunohistochemistry showed that these cells were Iba1 positive (Fig. 2C and D). Considering that we found monocytes in the peripheral blood to express Iba1, just like in other animal classes and humans (Imai et al., 1996) and found strong evidence that azurophils also express Iba1 (Dervas et al., 2023), we interpretated the above-mentioned cells as monocyte and azurophil precursors (i.e. promonocytes and proazurophils). In HE-stained sections, promonocytes were 8-10 μm in diameter, round cells with homogenous pale amphophilic to eosinophilic cytoplasm and a central to eccentric round nucleus. Proazurophils were of similar size, shape and with similar nuclear features, but had more intensely eosinophilic cytoplasm (Fig. 2C and D).

The ultrastructural examination of promonocytes confirmed their size. Their eccentric nucleus exhibited a moderate amount of peripheralized heterochromatin. The cytoplasm formed thin projections and contained abundant organelles, i.e. mitochondria, rough endoplasmic reticulum (RER) and lysosomes as well as a low to moderate number of small electron-dense granules (Supplemental Fig. 1D). Proazurophils were slightly smaller. Similar to monocytes, they carried organelles (mitochondria, rough endoplasmic reticulum) and small electron-dense granules in the cytoplasm, the latter less numerous than in promonocytes. They also exhibited variably sized (0.1-0.8 μm) vacuoles filled with finely granular material of low electron density, in variable numbers (Supplemental Fig. 2A). The round nucleus exhibited a moderate amount of clumped heterochromatin (Supplemental Fig. 2A). Some proazurophils also contained occasional 0.5 -2.5 μm irregularly shaped, lamellar, myelin-like material in the cytoplasm (Supplemental Fig. 2B).

<u>Lymphopoiesis.</u> Lymphoid cells could not be readily identified in HE stained sections, possibly due to their morphological similarity to erythropoietic precursors cells. Also, the immunohistochemical staining for the T cell marker CD3 did not identify individual or clusters of T cells. Attempts at the RNA-ISH for CD20 mRNA to identify B cells in the bone marrow were unsuccessful, likely due to RNA degradation during the decalcification process. However, TEM provided evidence of rare individual (pro)lymphocytes in the bone marrow, as it revealed small (3-4 µm in diameter) round cells dominated by a large central round nucleus with clumped heterochromatin that was surrounded by scant cytoplasm with a few mitochondria (Supplemental Fig. 2C).

<u>Plasma cells.</u> In HE-stained sections, cells with a morphology consistent with plasma cells in other animal classes were not observed. However, TEM revealed rare individual cells with plasma cell morphology in the bone marrow of one juvenile animal (B18). These cells were elongate to ovoid, 10-12 μm in diameter, with an eccentric round nucleus that contained centrally clumped heterochromatin (nucleolus) as well as heterochromatin that was radially arranged in the nuclear periphery (consistent with the “clockface appearance” of plasma cell nuclei in other animal classes (Bortnick and Murre, 2016)). The cytoplasm contained abundant rough endoplasmatic reticulum that formed closely spaced cisternae, a high number of mitochondria and a Golgi apparatus (Supplemental Fig. 2D).

#### 3.3.2 Bone marrow in BIBD-positive animals

In BIBD-positive animals, the bone marrow had a principally identical composition. However, most cell types were found to be reptarenavirus infected, as they carried the pathognomonic intracytoplasmic IBs (Fig. 3A-C). These were most prominent in proerythrocytes where they presented as clearly demarcated cytoplasmic IBs in HE-stained sections and reptarenavirus nucleoprotein (NP) accumulations in immunostained slides; these cells were concentrated around the sinus venosus (Fig. 3A). In addition, a few granulopoietic and monocytopoietic cells (progranulocytes, proazurophils and promonocytes, respectively) exhibited obvious IBs in HE-stained sections and distinct focal intracytoplasmic viral NP expression (Fig. 3C). The majority of the remaining cells lacked obvious IB but showed viral NP expression (Fig. 3B and C) which was generally evident as diffuse, occasionally granular cytoplasmic staining (Fig. 3C). Cells with viral antigen expression and IBs were also found in the lumen of the sinus venosus, indicating the release of infected cells into the blood stream (Fig. 3B and C). Further evidence of the latter was obtained from the buffy coat in which the ultrastructural examination highlighted intracytoplasmic IBs in abundant erythrocytes and numerous leukocytes with monocyte morphology (Fig. 3D).

**Fig. 3.**
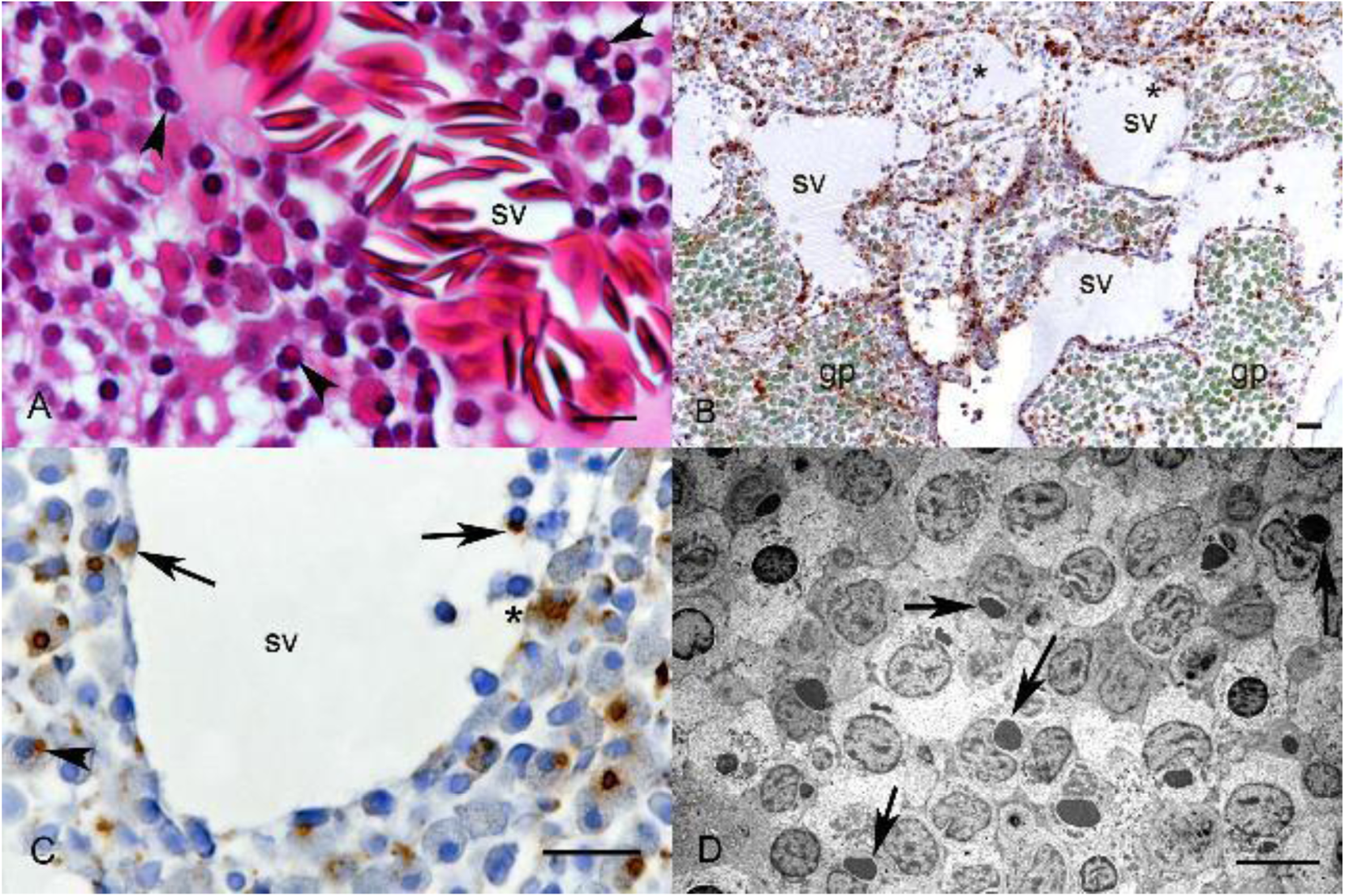
The bone marrow in BIBD. **A. Inclusion bodies in proerythrocytes** (animal A3, 2 years). HE-stained section, showing eosinophilic cytoplasmic inclusion bodies (IB) mainly in proerythrocytes (arrowheads), these form dense aggregates around the sinus venosus (sv). HE stain, bar = 20 µm. **B. Reptarenavirus infects a large proportion of haematopoietic cells.** BIBD positive animal (A3, 8 years) showing extensive viral nucleoprotein (NP) expression in haematopoietic cells. The asterisk indicates infected cells in the lumen of the sinus venosus (sv). gp: granulopoietic cells, as indicated by the green colouration of their cytoplasmic granules. Immunohistochemistry, haematoxylin counterstain, bar = 20 µm. **C. Inclusion body formation occurs in different types of haematopoietic cells** (animal A4, 2 years). IBs are present in proerythrocytes around and in the lumen of the sinus venosus (sv, arrows) and in cells with progranulocytic appearance (arrowhead). A more diffuse cytoplasmic NP expression is seen in a cell with progranulocyte morphology (asterisk). Immunohistochemistry, haematoxylin counterstain, bar = 20 µm. **D. Inclusion bodies in cells in the peripheral blood.** Buffy coat of a BIBD positive boa (animal A12, 2 years). Numerous cells with monocyte morphology show characteristic intracytoplasmic IBs, identified as juxtanuclear aggregates of electron dense material, i.e. viral nucleoprotein (arrows). The affected cells are identified as monocytes based on their high amount of cytoplasm rich in organelles, and the eccentric nucleus. TEM, bar = 5 µm.

### 3.3 Thymus

#### 3.3.1. Location, morphological features and composition

Grossly, the thymus presented as an approximately 25 x 5 x 5 mm unilobed organ of light brown colour located adjacent to the trachea and cranially to the thyroid gland, in close proximity to carotid artery, jugular vein, and vagal nerve (Fig. 4A). It was evident in all animals regardless of age, without obvious evidence of a reduction in size, i.e. involution.

**Fig 4.**
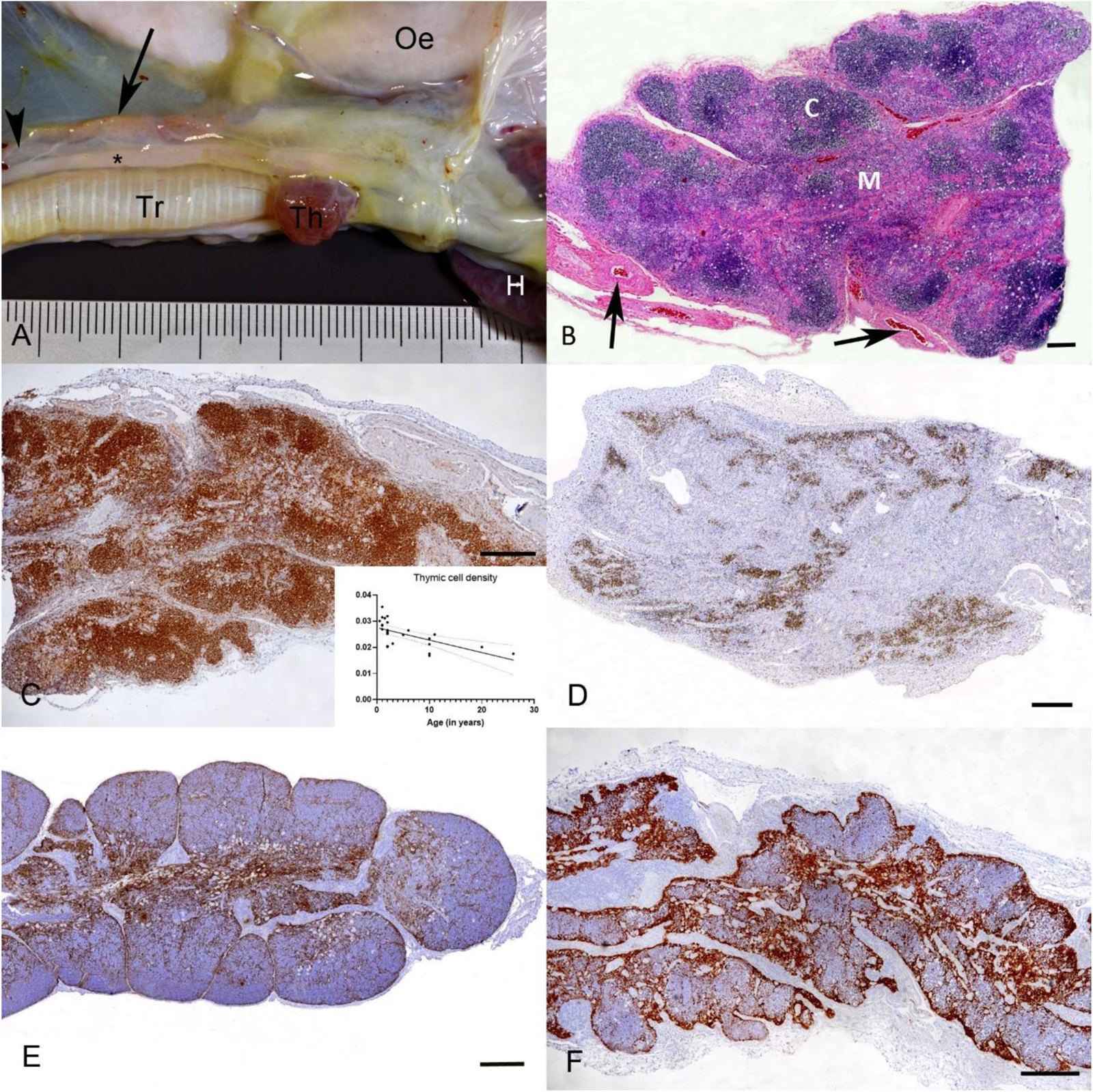
Anatomic features of the thymus in *B. constrictor*. **A. Thymus of an adult snake**, situs (animal B7, 10 years). The thymus (arrow) is a unilobed organ of light brown colour adjacent to the trachea (Tr), cranial to the thyroid gland (Th), and close to the carotid artery (asterisk) and vagus nerve (arrowhead). Oe: oesophagus, H: heart. **B. Histological overview of the thymus of an adult snake** (animal B8, 6 years). Cortex (C) and medulla (M) are moderately distinct. The cortex is composed of several small lobes that are rich in small lymphocytes. Small to medium-sized blood vessels are found within the capsule (arrows). HE stain, bar = 100 μm. **C-F. Composition of the thymus in young and older animals. C. T cells in the thymus of a young animal** (B3, 1 year). Staining for CD3, a T cell marker, highlights the prominent cortex comprised of small T cells that form cell rich aggregates. Inset**: Thymic overall cell density**. The morphometric analysis of the total number of nuclei (counted on HE-stained sections) revealed a low negative correlation of the thymic cell density with age (p<0.001). **D. T cells in the thymus in an old animal** (B7, 8 years). The cellularity of the cortex is reduced due to a markedly lower number of thymocytes (small lymphocytes, CD3+). **E. Epithelial cells in the thymus of a young animal** (D1, 1.5 years). Epithelial cells (pan-cytokeratin+) are mainly found in the medulla, in low to moderate numbers. **F. Epithelial cells in the thymus of an old animal** (B7, 10 years). The medulla is prominent due to the high number of epithelial cells (pan-cytokeratin+; epithelial hyperplasia), whereas the cortex is markedly reduced. Immunohistochemistry, haematoxylin counterstain. Bars = 250 μm.

Histologically, the thymus was composed of several small lobes (Fig. 4B), each divided into smaller lobules by trabeculae that extended from a dense fibrous capsule into the cortex and consisted of collagen as well as elastic and reticulin fibres (Fig. 4B).

There was evidence of reducing cellularity with age. This was confirmed by the morphometric analysis performed on HE-stained sections which revealed a negative correlation of the total nuclear count (total number of cells) with age (p<0.001; R2=0.371, F=14.18) (Fig. 4C, inset).

Also, in younger animals, cortex and medulla were clearly discernible; these were less obvious in older individuals. In young animals, the cortex was supported by a thin meshwork of interlacing, cytokeratin positive epithelial-reticular cells and reticulin fibres. Embedded into this were abundant T cells (CD3+; thymocytes). The lymphocytes occasionally formed follicle-like aggregates. Regardless of the animals’ age, a moderate number of macrophages (Iba1+) were also present in the cortex and/or at the corticomedullary junction. Heterophils were mostly found in low, rarely in moderate to high numbers in the outer cortex, at the transition to the fibrous capsule. In older animals (approximately >6 years of age), the overall number of lymphocytes (T cells) in the cortex was lower (Fig. 4 C and D) and. the epithelial component was more prominent (Fig. 4E and F) (See also 3.3.2.).

The medulla contained abundant disseminated epithelial cells (cytokeratin+) and scattered large round acidophilic cells with a small dark nucleus, morphologically consistent with myoid cells (Varga et al., 2019) (Fig.5A). The ultrastructural examination showed that the epithelial cells varied in size (ranging from 10-12 μm in diameter) and shape (cuboidal to polygonal). In some animals, epithelial cells were found to form multifocal ductular structures (Fig. 5B). Epithelial cells within the ductular structures often displayed cilia, characterized by a dense brush border of cilia axonemes supported by basal bodies along the apical plasma membrane, and a basally located round nucleus (Fig. 5C). Ultrastructurally, myoid cells had an euchromatic nucleus and contained regular myofibrils that were arranged in parallel arrays in the cytoplasm (Fig. 5B and D). In addition, both cortex and medulla contained scattered random Ig Y and Ig M expressing cells, respectively, CD20 positive B cells (Fig. 5E and F) and single cells with ultrastructural features consistent with plasma cells (prominent Golgi complex, abundant mitochondria and a nucleus with marginalized crumpled heterochromatin). Rare heterophils were also found in cortex and medulla; they were occasionally arranged perivascularly.

**Fig. 5.**
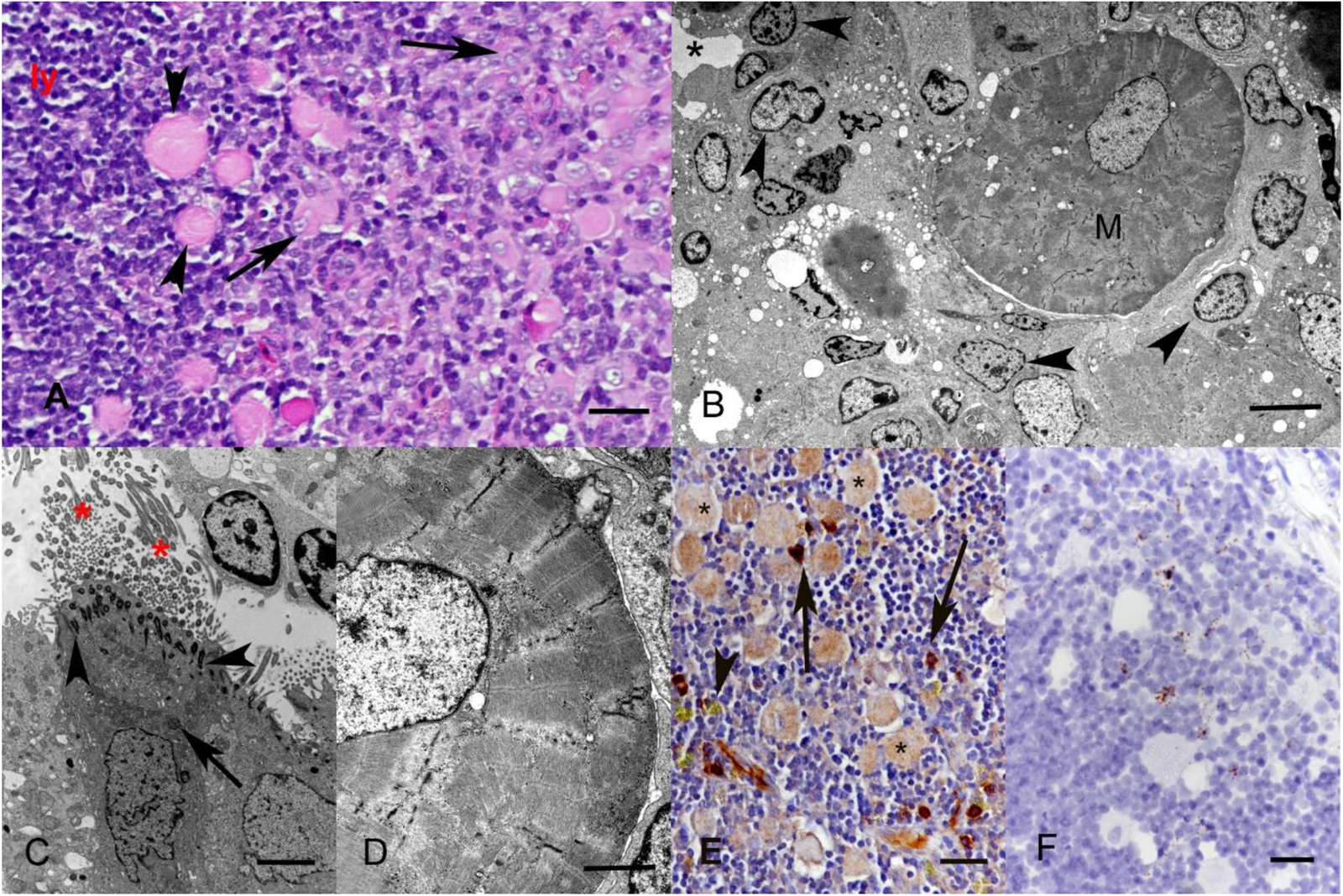
Microarchitecture and ultrastructure of the thymus in *B. constrictor*. **A.** Histological features of the medulla (animal D2, 1.5 years). There are numerous epithelial cells (arrows) and scattered large myoid cells (arrowheads), ly: lymphocytes (thymocytes). HE stain, bar = 20 µm **B. Ultrastructural features of the medulla** (animal A16, 3 months). The centre is occupied by a myoid cell (M) surrounded by epithelial cells (arrowheads). Epithelial cells also form duct-like structures (asterisk: duct lumen). TEM, bar = 10 µm. **C. Ductal epithelial cell** (animal A16, 3 months) with apical ciliation (asterisks), characterized by a brush border of cilia axonemes supported by basal bodies (arrowhead). The roundish nucleus is basally located, euchromatic, with marginal condensation of chromatin. The cytoplasm shows small vacuoles and a high number of mitochondria (arrow). TEM, bar = 2 µm. **D. Myoid cell** (animal A16, 3 months) with an euchromatic nucleus and transverse myofibrils arranged parallelly in a regular pattern. TEM, bar = 2 µm. **E. Ig Y+ cells** (animal B10, 1 year) are found multifocally in the medulla (arrows). The arrowhead points at a heterophil which can be recognised due to the green colouration of its granules after any immunohistochemical stain using DAB as chromogen (arrowhead). Asterisk: myoid cells. Immunohistochemistry, haematoxylin counterstain, bar = 10 µm. **F. CD 20 mRNA expressing (B) cells** (animal B8, 6 years) are present in low numbers scattered throughout the thymus. RNA-ISH, haematoxylin counterstain, bar = 10 µm.

#### 3.3.2. Age-related and incidental histopathological changes

The thymus exhibited several age-related and/or incidental changes: epithelial or thymic cysts, and epithelial hyperplasia. Epithelial cysts (Pease, 1956) were present in the medulla of the majority of animals (35/44, 80%), regardless of age, sex or BIBD status. Histologically, they were represented by variably sized cystic spaces filled with pale eosinophilic to basophilic homogenous fluid and lined by a single layer of flattened to columnar epithelium. Some epithelial cells showed evidence of intracytoplasmic mucin (goblet cell transformation) (Fig 6A). Two animals (B15, BIBD-positive; B4, BIBD and reptarenavirus negative) showed histologic evidence of multiple thymic cysts (Strollo and Rosado-de-Christenson, 2012). These were noticeably larger than epithelial cysts (up to 1 cm in diameter) and contained a large amount of pale eosinophilic fluid and sloughed degenerate cells. The cysts were lined by a single-to multilayered epithelium with occasional ciliated cells. Older individuals with epithelial cysts, both BIBD-positive and -negative, exhibited an increased number of cuboidal to columnar epithelial cells (cytokeratin+) in the medulla. Cells were often forming tubules or cords that were frequently dilated (Fig. 6B).

**Fig. 6.**
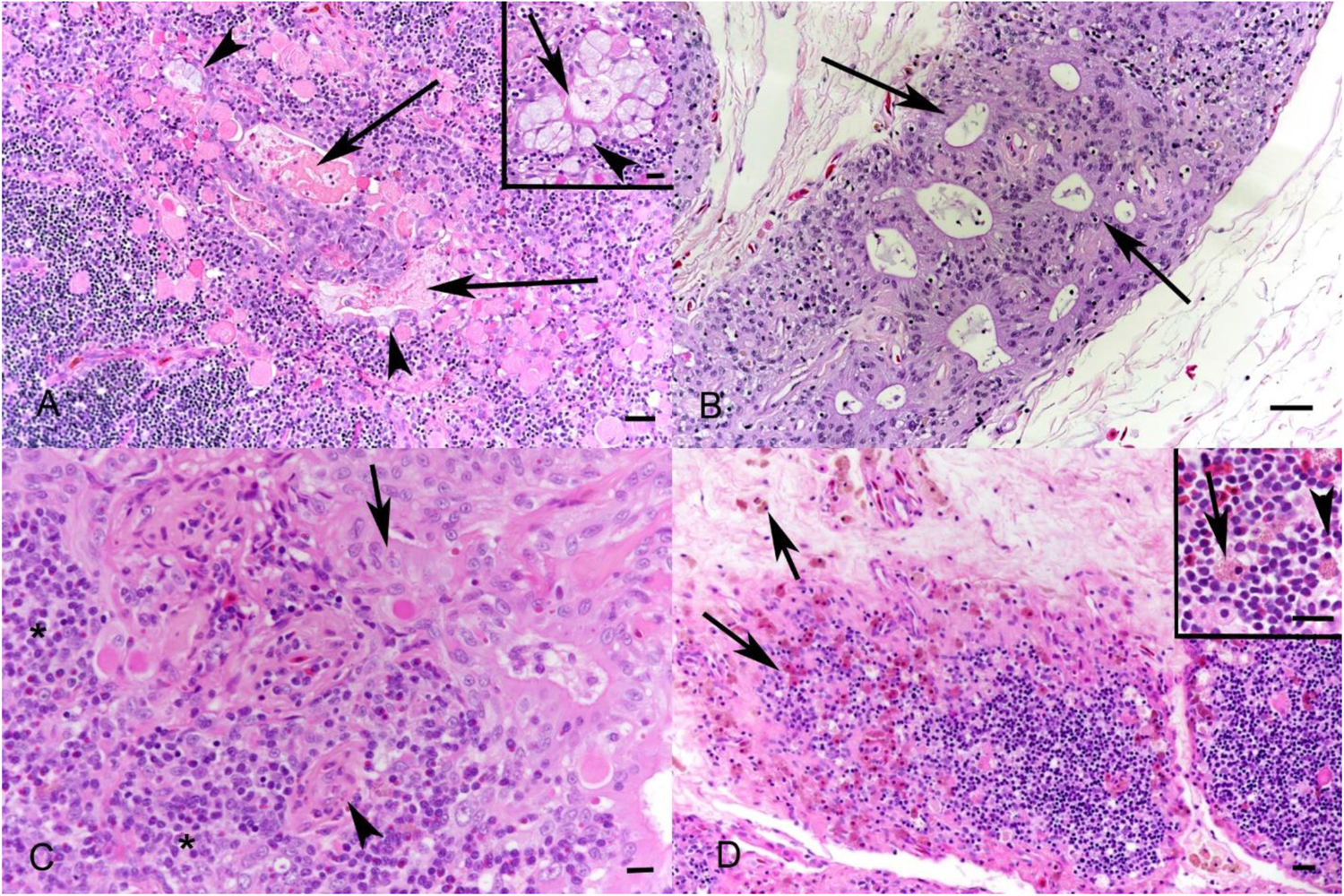
Histological changes in the thymus related to age and BIBD/reptarenavirus infection. **A.** Epithelial cysts (animal B4, 8 years , BIBD and reptarenavirus negative). The medulla exhibits cystic spaces that are filled with pale eosinophilic homogenous material (arrows). The epithelium occasionally shows foamy and distended cytoplasm (arrowhead), consistent with goblet cell transformation. Inset: Higher magnification of goblet cell transformation (arrowhead). The cells also exhibit apical ciliation (arrow). HE stain, bars = 20 μm and 10 μm (inset). **B. Epithelial hyperplasia** (animal A2, 1 year, BIBD positive). Medulla with columnar epithelial cells that form dilated tubules (arrows). HE stain, bar = 20 μm. **C, D. Histological changes in the thymus of snakes with BIBD**. **C. Inclusion body formation** (animal A21, 11 years) in the cytoplasm of medullary epithelial cells (arrow), vascular endothelial cells (arrowhead) and cortical lymphocytes (asterisks). **D. Infiltrating ”granular cells”** (animal A11, 2 years). Marked infiltration of capsule, cortex and medulla by often perivascular “granular cells” (arrows). The cytoplasm of these cells is filled with a high number of yellow to brownish granules (inset). HE stain, bars = 20 μm and 10 μm (inset).

#### 3.3.3. The thymus in BIBD-positive animals

BIBD-positive animals showed intracytoplasmic IBs and reptarenavirus NP expression in thymocytes, epithelial and vascular endothelial cells (Fig. 6C). They also exhibited numerous round cells (8-12 μm in diameter) with abundant cytoplasmic yellow granules that were negative for Fe2+. The cells did not express any of the tested markers and were hence tentatively called “granular cells” (Fig. 6D).

The morphometric analysis, performed on the thymus of 10 BIBD/reptarenavirus-negative and 14 BIBD-/ reptarenavirus-positive animals, did not reveal any significant differences in total cell number, number of apoptotic cells (cleaved caspase-3+) and number of T cells (CD3+) between BIBD-positive and BIBD-negative animals (Supplemental Table 2), indicating that the disease does not affect composition or cell turnover and survival in the spleen.

### 3.3 Spleen

#### 3.3.1. Location, morphological features and composition

The spleen is a spherical organ located cranially to the pancreas and the gall bladder, at the transition of the pyloric region of the stomach to the duodenum (Fig. 7A), delineated by a thick fibrous capsule from which fibrous connective tissue cords stretched towards and divided the parenchyma into moderately demarcated lobules (Fig. 7B). The serosal surface was covered by a mesothelial cell layer. Beneath the capsule was a thick layer of elastic fibers, broad collagen fibers and smooth muscle cells (α-SMA+), all present at approximately equal amounts and also forming the septae. Both capsule and septae contained numerous large arteries and veins (Fig. 7C and D).

**Fig. 7.**
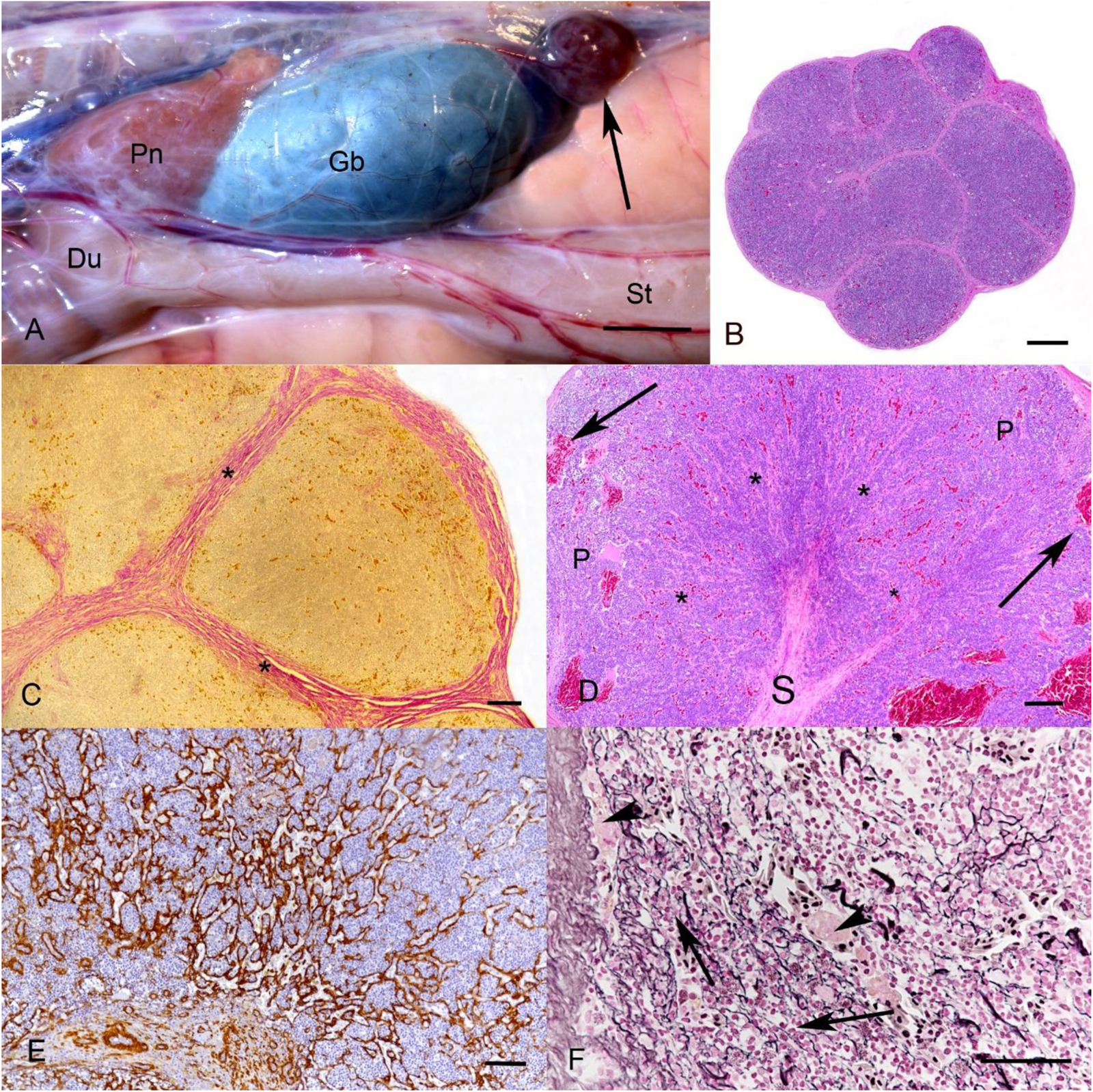
Anatomic features of the spleen in *B. constrictor*. **A.** Spleen, situs (animal B2, 8 years). The spleen (arrow) is located cranially to the pancreas (Pn) and the gall bladder (Gb), at the transition of the pyloric region of the stomach (St) to the duodenum (Du). **B, C. Cross section of a spleen highlighting the capsule and septal tissue** (B7, 10 years). **B.** The spleen is surrounded by a fibrous capsule from which fibrous connective tissue cords stretch and divide the parenchyma into moderately demarcated lobules. HE stain, bar = 100 μm. **C.** The fibrous connective tissue cords consist of thick, parallelly arranged collagen fibres (asterisks). Van Giesson stain, bar = 200 μm. **D. Transition of the capsule-septal tissue to the perilymphoid fibrous zone (PLFZ)** (animal A3, 2 years). The PFLZ (asterisks) presents as an intermediate zone between the septal connective tissue stalk (S) and the white pulp (P). Note the presence of dilated blood-filled veins in the pulp (arrows). HE stain, bar = 200 μm. **E+F. PFLZ** (animal B7, 10 years). **E.** The PFLZ consists of a meshwork of small to medium sized veins that are supported by α-SMA positive reticular cells. Immunohistochemistry, haematoxylin counterstain, bar = 200 μm. **F.** Reticulin fibres surround small veins (arrowheads) and the lymphoid tissue (arrows). Reticulin stain, bar = 200 μm.

The parenchyma was comprised of lymphoid tissue (“white pulp”) and a perilymphoid fibrous zone (PLFZ), as described for other reptile species (Tanaka and Hirahara, 1995). The PFLZ presented as an intermediate zone located between the capsule-septal tissue and the lymphoid tissue and consisted of a meshwork of small to medium sized veins that were supported by reticulin fibres and α-SMA positive reticular cells (Cheng et al., 2019) (Fig. 7E). While a clear distinction between PLFZ and lymphoid tissue was not possible in HE-stained sections, as the border of the PLFZ was obscured by lymphocytes, it was apparent in the reticulin stained sections and in sections stained for α-SMA (Fig. 7E and F).

The white pulp was composed of abundant lymphocytes, arranged in sheets in the centre of each lobule. T cells (CD3+) made up 90-95% of the lymphocytes (Fig. 8A), while B cells (CD20 mRNA+) did not amount to more than 5%, maximally 10%. The T cells were occasionally arranged in concentric aggregates reminiscent of lymphatic follicles in other species, with a central round lighter zone resembling a germinal centre (Fig. 8A). In contrast, B cells (CD 20 mRNA+) were generally randomly distributed individual cells that were rarely arranged as small, random aggregates of 5-10 cells (Fig. 8B). In HE-stained sections, there was no evidence of cells with plasma cell morphology as it is known from other animal classes (Bortnick and Murre, 2016). However, the ultrastructural examination revealed occasional cells with plasma cell features (Fig. 8C). In addition, a low to moderate number of IgY and IgM expressing cells were present. These were generally found scattered in the white pulp, either as random individual cells or, more frequently, in close proximity to the PFLZ (Fig. 8D, E). Morphological compartmentalisation, with red pulp and/or evidence of periarteriolar lymphoid sheath (PALS)/peri-ellipsoid lymphocyte sheath (PELS) structures was not evident.

**Fig. 8.**
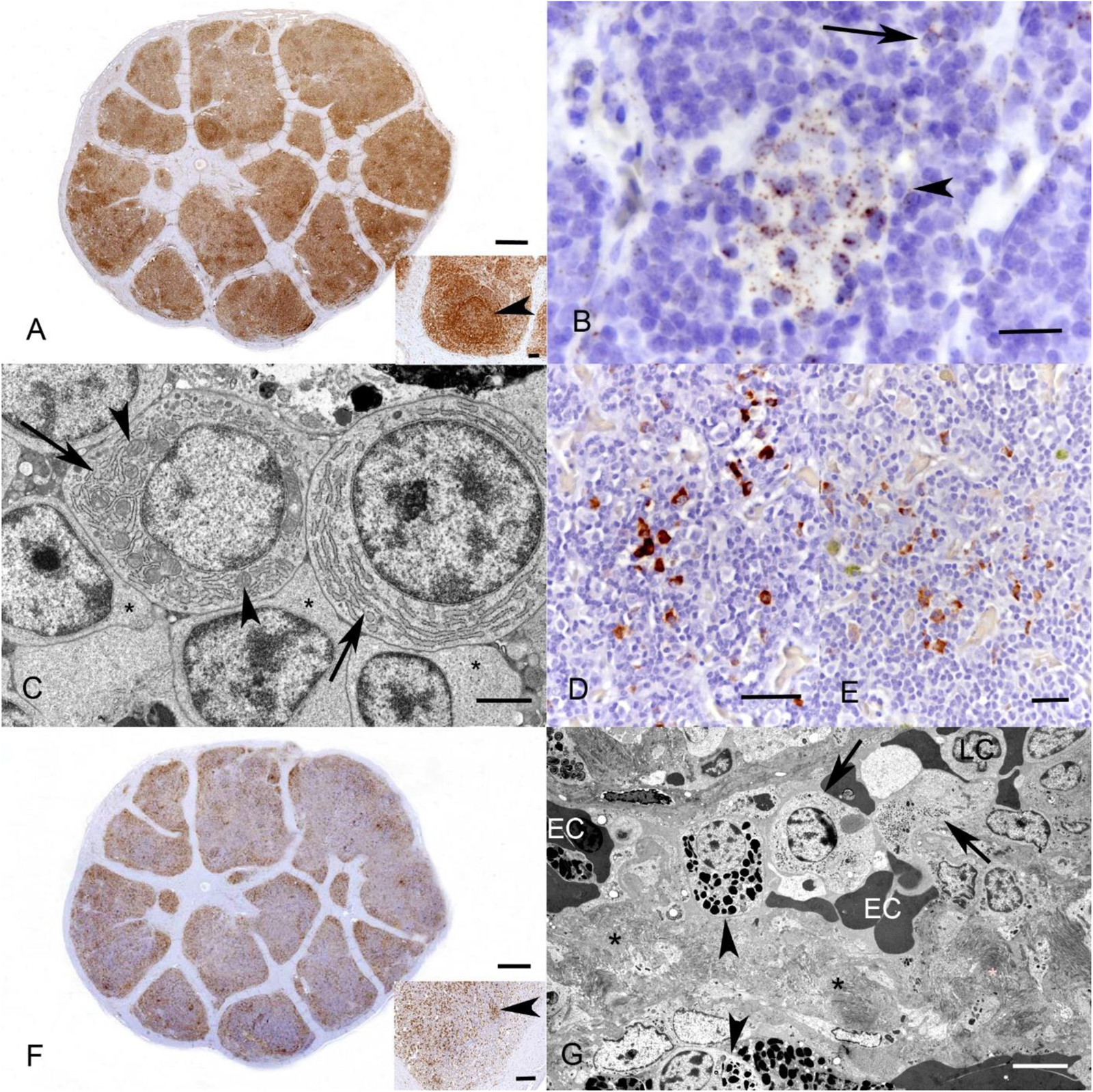
Splenic cell populations. **A. T cells** (animal B7, 10 years). Staining for the T cell marker CD3 shows that the white pulp consists primarily of T cells arranged in sheets in the centre of each lobule, occasionally arranged in a follicle-like manner (inset: arrowhead). Immunohistochemistry, haematoxylin counterstain, bar =100 μm**. B. B cells** (animal A3, 2 years). Cells positive for CD20 mRNA (i.e. B cells) are present at low numbers, most frequently isolated (arrow) and occasionally forming small aggregates (arrowhead). RNA-ISH, haematoxylin counterstain, bar = 20 μm**. C. Plasma cells** (animal E, adult), identified based on their ultrastructural features, i.e. a prominent Golgi complex (arrowheads) and abundant mitochondria (arrows) in the cytoplasm and a nucleus with marginalized crumpled heterochromatin. Adjacent to the plasma cells are lymphocytes (asterisks). TEM, bar = 2 μm**. D, E. Ig-expressing cells** (animal B6, 10 years). **D.** Several individual IgY-positive cells in the spleen. **E.** Several individual IgM-positive cells in the spleen. Immunohistochemistry, haematoxylin counterstain, bars = 20 μm. **F. Monocytes/macrophages** (animal B7, 10 years) are present at a moderate number in the white pulp. They are predominantly dispersed throughout the organ and rarely form small aggregates (inset: arrowhead). Immunohistochemistry (Iba1), haematoxylin counterstain, bar=100 μm. **G. Ultrastructural features of the splenic cell populations** (animal E, adult). Macrophages (arrows) with an eccentric nucleus and abundant cytoplasm with a high number of organelles, and heterophils (arrowheads), with abundant intracytoplasmic electron dense granules between dense connective tissue fibrils (asterisks) and few erythrocytes (EC). TEM, bar = 4 μm.

In most cases, macrophages (Iba1+) were seen in moderate to high numbers interspersed between lymphocytes in the white pulp and in the PFLZ; they occasionally formed small aggregates (Fig. 8F (inset) and F).

In the majority of animals two further, noticeably less abundant cell populations were seen. The first population were Iba1 positive haemosiderin-laden (iron stain: positive) macrophages (i.e. siderophages) that were often located around small vessels, within the PFLZ and/or adjacent to the capsule. The second population was very small and represented by round cells of the same size as siderophages but with a moderate amount of cytoplasm containing numerous yellow granules (identical to the granular cells observed in the thymus of BIBD-positive animals, and like these negative for Fe2+ and for the expression of CD3, IgY, IgM and Iba1; they were hence also tentatively called “granular cells”.

#### 3.3.2. Histopathological changes in BIBD- and reptarenavirus-negative animals

In all but one BIBD-negative animal, the spleen did not exhibit any pathological changes. One snake (B9) showed a single, well circumscribed granuloma in the spleen. It replaced part of the white pulp and was characterized by a central area of necrosis, surrounded by a rim of macrophages and peripheral fibroblasts. The Ziehl-Neelsen stain did not identify any acid-fast bacteria in the lesion. The animal did not exhibit any further inflammatory changes in any other inner organs.

#### 3.3.2. The spleen in BIBD-positive animals

The spleens of BIBD-positive snakes appeared overall more cell rich than those of control animals; this was confirmed by the morphometric analysis which revealed a significantly higher overall cellularity in BIBD-positive snakes (mean=0.022, CI=0.019-0.025) than BIBD-negative animals (mean=0.028, CI=0.026-0.029), with t= -3.71, df=25, p<0.01 (Fig.9A). However, this phenomenon was neither associated with a selective increase in T cells (CD3+) or macrophages (Iba1+) nor with a difference in the extent of cellular turnover (assessed based on the number of cleaved caspase 3+ apoptotic cells). The results of the morphometric analysis are provided in Supplemental Table 2.

**Fig. 9.**
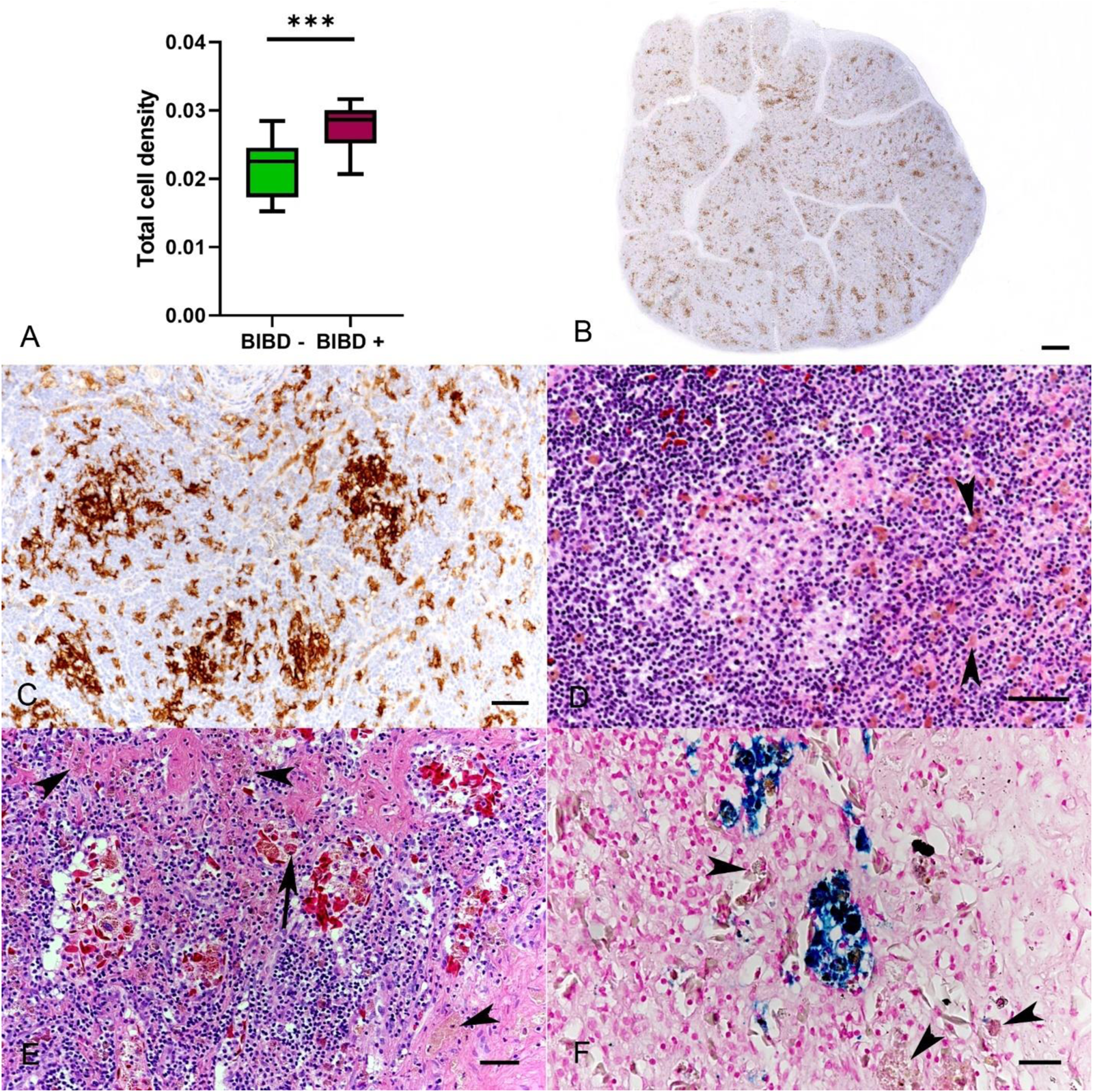
Histological changes in the spleen of BIBD positive animals. **A. Higher overall cell density in the spleen of snakes with BIBD**. The morphometric analysis of the total number of nuclei (counted on HE-stained sections from the spleens of 17 BIBD positive snakes and 10 BIBD and reptarenavirus negative snakes) revealed significantly higher overall cellularity in BIBD (p<0.01). **B, C. Monocyte/macrophage rearrangement. B.** Animal A9, 2 years. Staining for Iba1 shows multiple disseminated, variably sized focal macrophage aggregates. **C.** Animal A13, 8 years. Macrophages (Iba1+) are present as individual cells and, more frequently, arranged in small groups. Immunohistochemistry, haematoxylin counterstain, bars = 100 μm (B) and 20 μm (C). **D.** Animal A20, 6 years. In the HE-stained section, macrophage aggregates are evident as groups of large cells with foamy to vacuolated cytoplasm. Several “granular cells” (arrowheads) are also present. HE stain, bar = 20 μm (inset). **E, F. Siderophages and “granular cells”** (animal B7, 8 years). **E.** Haemosiderin-laden macrophages with dark granular cytoplasm (arrow) and “granular cells” with their abundant yellowish cytoplasmic granules (arrowheads). HE stain, bar = 20 μm. **F.** Siderophages are identified as they contain Fe2+ in their cytoplasm (blue deposits). The adjacent “granular cells” are negative (arrowheads). Prussian blue stain, bar = 20 μm.

Nonetheless, there was a noticeable difference in the splenic arrangement of macrophages (Iba1+). While the Iba1 positive cell population was more or less evenly dispersed in the spleen of the control animals (Fig. 8B), it BIBD-positive often formed aggregates, indicating monocyte/macrophage rearrangement consistent with a granulomatous response, in BIBD-positive snakes (Fig. 9B, C). In addition, both siderophages (consistent with haemosiderosis) and “granular cells” appeared overall noticeably more abundant (Fig. 9D-F). Both cell populations were seen as individual cells or as aggregates of up to 15 cells; they were also found in the lumen of vessels (Fig. 9E and F), indicating increased local phagocytic and digestive activity.

The spleens of BIBD-positive animals also contained more heterophils than the spleens of control animals. These were mainly located in the periphery of the lobules, in the PFLZ, and/or at the transition to the splenic capsule. In eight BIBD-positive animals (A3, A5, A14, A19-A22) variably sized, thin-walled vascular structures were noted within the PFLZ; these were filled with pale eosinophilic homogenous material and lined by a single layered endothelium (interpreted as dilated lymphatic vessels; lymphangiectasia) (Fig.10A). Another three BIBD-positive animals (A6, A8, A9) exhibited well demarcated granulomas (granulomatous splenitis, Fig. 10B). Also in these cases, the Ziehl-Neelsen stain did not reveal any acid-fast bacteria in the lesions. The only concurrent lesion seen in the inner organs of these three animals was a variable degree of a heterophilic colitis.

**Fig. 10.**
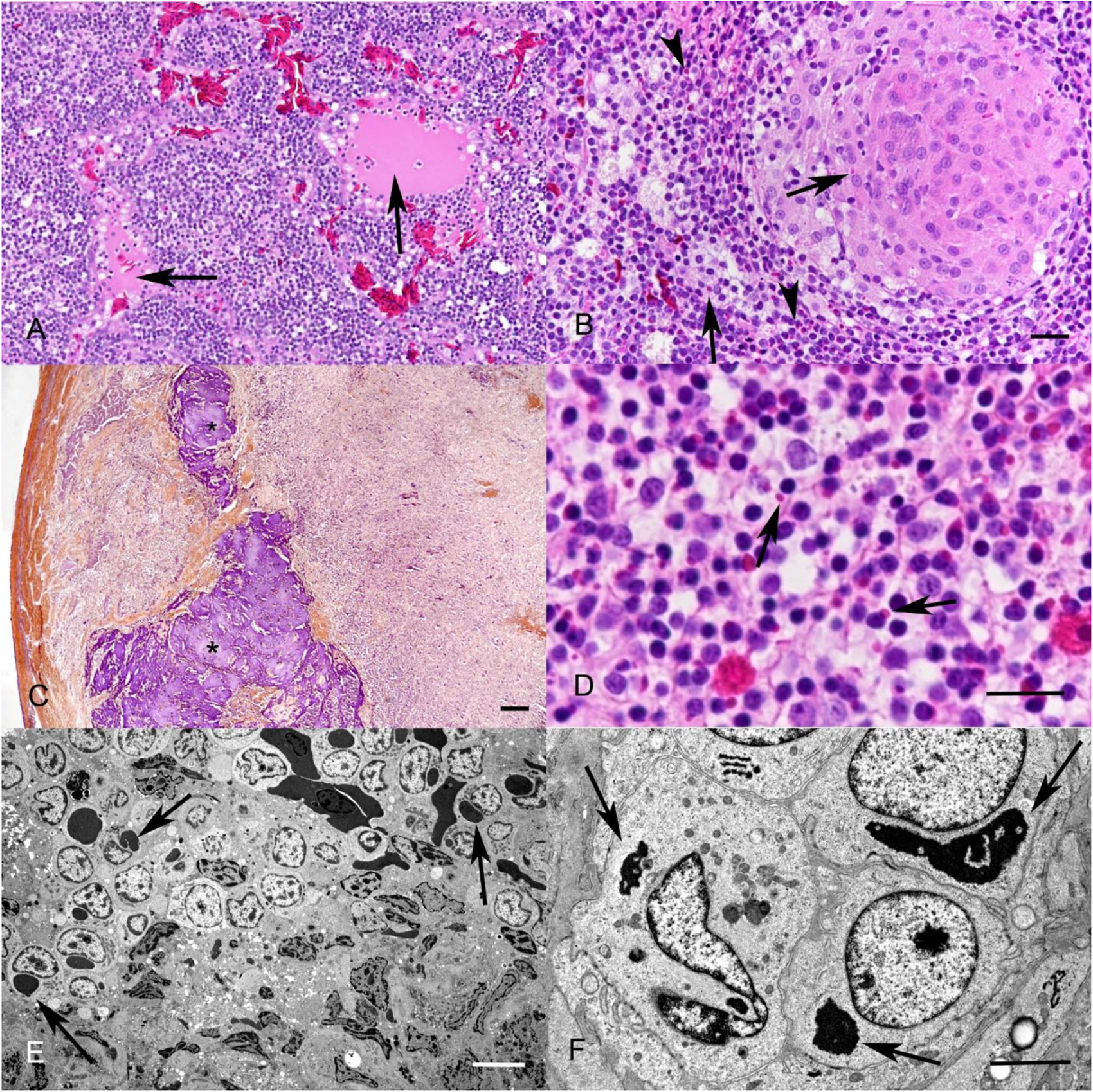
Histological changes in the spleen of BIBD positive animals. **A. Lymphangiectasia** (animal A13, 8 years). Variably sized, thin-walled vascular structures filled with pale eosinophilic homogenous material (arrows), HE stain, bar = 10 μm **B. Granulomatous splenitis** (animal A9, 2 years). Granuloma with inclusion bodies (IB) in macrophages (arrow). In the white pulp in the periphery of the granuloma, scattered lymphocytes (arrowheads) and macrophages (arrow) with IB are seen. HE stain, bars =10. **C. Fibrin deposition** (animal A20, 6 years). Multifocal fibrinous deposition (asterisks). PTAH stain, bar=100 μm. **D-F. Inclusion body formation. D.** Animal A20, 6 years. IB in lymphoid cells (arrows). Bar= 20 μm. **E+F. Ultrastructural features of splenic cells.** Animal E, adult. Round cells with the morphology of macrophages (with abundant cytoplasm rich in organelles and an eccentric nucleus. Many of these exhibit intracytoplasmic inclusion bodies, identified as juxtanuclear electron dense material (arrows). TEM, bar = 10 μm. **F.** Animal E, adult. Deposition of juxtanuclear electron dense material of variable size in splenic cells, e.g. lymphocytes (bottom and upper right). TEM, bar = 2 μm.

Two BIBD-positive snakes (A20, B16) showed large multifocal fibrin deposits in the splenic parenchyma and/or the lumen of splenic vessels (Fig. 10C). In both animals, concurrent changes in other organs were seen, namely a mild chronic interstitial pneumonia and a heterophilic nephritis and/or colitis, indicating a septicaemic process. The latter could not be confirmed as a bacteriological examination was not initiated at the time of necropsy.

In addition, the spleens contained abundant lymphocytes and macrophages with intracytoplasmic IB and reptarenavirus NP expression (Fig. 10D-F) in all animals.

### 3.4 Lymphatic tissue of the alimentary tract

#### 3.4.1. Location, morphological features and composition

<u>Oesophageal tonsils</u>. These presented grossly as multiple circumscribed whitish to dark red, 1-2 mm in diameter luminal mucosal protrusions along the entire length of the oesophagus (Fig. 11A). Histologically, these corresponded to large, ellipsoid, cell dense, raised aggregates of T cells (CD3+) and a few B cells (CD 20 mRNA+) arranged around numerous capillaries (Fig. 11B, C). Scattered individual cells expressing IgM and IgY were also noted. The overlying epithelium contained multiple intraepithelial nests of T cells and some macrophages (Iba1+) (Fig. 11C).

**Fig. 11.**
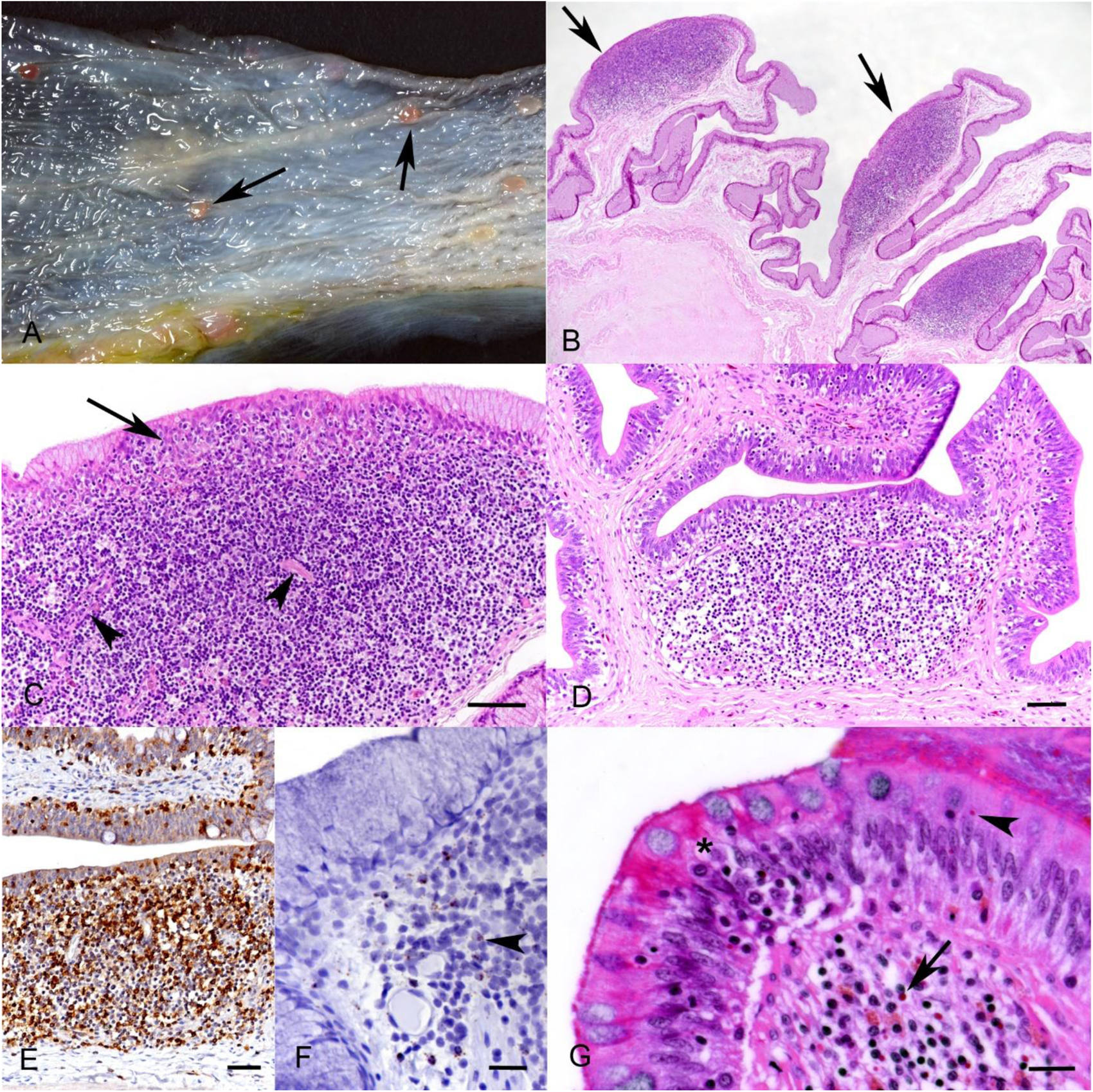
Lymphatic tissue of the alimentary tract. **A-C. Oesophageal tonsils** (animal B10, 7 years)**. A. Gross image, mucosal surface of a longitudinally opened oesophagus.** The oesophageal tonsils present as multiple raised whitish to red nodular mucosal protrusions (arrows). **B, C. Histological features**. **B.** Overview, showing large, ellipsoid, protruding dense aggregates of lymphoid cells in the submucosa (arrows). **C. Closer view** of the top left lymphoid aggregate in B, highlighting the dense aggregate of mostly lymphocytes, with embedded capillaries (arrowheads). The overlying squamous epithelium contains small nests of lymphocytes and macrophages (arrow). HE stains, bars = 100 μm and 50 μm. **D-F. Gut associated lymphoid tissue (GALT), small intestine** (animal C3, 1 year)**. D.** Nodular, well demarcated submucosal aggregate of lymphocytes and fewer macrophages. HE stain, bar = 50 μm. **E. T cells.** The GALT is mainly comprised of T cells; these are also infiltrating the overlying epithelium. Immunohistochemistry, haematoxylin counterstain, bar = 50 μm. **F. B cells** (CD20 mRNA+) are rare and mainly located in the periphery of the lymphocyte aggregates (arrowhead). RNA-ISH, haematoxylin counterstain, bar = 20 μm. **G**: Animal A20, 6 years. IB formation in the mucosal epithelium (arrowhead) and in inflammatory cells (arrow). Note also the lymphocytes infiltrating the epithelial layer (asterisk). HE stain, bar = 20 μm.

<u>Gut associated lymphatic tissue (GALT)</u>. GALT was not grossly apparent but was histologically evident as small nodular, well demarcated, aggregates of round cells that occasionally expanded the wall in the small and large intestine (Fig. 11 D). It was predominantly comprised of T cells (CD3+), admixed with fewer macrophages (Iba1+), B cells (CD20 mRNA+), IgM and IgY expressing cells and heterophils (Fig. 11E, F).

#### 3.3.2 Changes in BIBD-positive animals

Intracytoplasmic IB and reptarenavirus NP expression were seen in abundant lymphocytes and macrophages of the GALT and oesophageal tonsils as well as in the epithelial cells of the overlying mucosa (Fig.11G). The majority of the BIBD-positive animals presented some degree of diffuse heterophilic infiltration of the Lamina propria and submucosa in the small and/or the large intestine (heterophilic enterocolitis). In these cases, the GALT was enlarged and contained more heterophils, predominantly in the periphery of the lymphoid aggregates. Additionally, in two BIBD-positive animals (A16 and A17), the stomach exhibited submucosal lymphatic tissue, presenting as small, well demarcated follicle-like accumulations of T cells (CD3+), admixed with rare individual B cells (CD20 mRNA+) and/or macrophages.

## 4. Discussion

The present study focused on the morphologic characterization of the haemolymphatic tissues of the *B. constrictor*, in order to gather reference data for its assessment under disease conditions. The *B. constrictor* is one of the globally most common snake species kept, traded and bred in captivity, and the main species affected by BIBD. Over the last decades, BIBD has been the most important viral disease of captive boid snakes worldwide. It has a detrimental effect on the affected individual and on entire breeding collections, hence it is concerning that its pathogenesis and its effect on the host’s immune response has so far not been elucidated (Chang and Jacobson, 2010; Schumacher et al., 1994). From an evolutionary perspective, reptiles, as the only living ectothermic amniotes, represent a link between ectothermic anamniotic fishes/amphibians and the endothermic amniotic birds/mammals. Therefore, a better understanding of their immune system may uncover important clues regarding the evolutionary history of the immune responses in vertebrates in general and could contribute to the conservation of reptile species. Here we provide detailed information on the anatomic features and cellular composition of bone marrow, thymus, spleen and lymphatic tissue of the alimentary tract in healthy *B. constrictor*, highlighting also age-associated differences. The comparative assessment of these organs in BIBD and reptarenavirus free snakes and those with the disease provides first insight into the immune response to or possible effects of reptarenavirus infection.

### 4.1. The haematopoietic tissues in B. constrictor in comparison to their equivalents in other animal classes

Our results in control animals show that the haemolymphatic tissues in *B. constrictor* have many similarities to their equivalents in other animal classes but there are also noticeable differences.

<u>Thymus</u>. In boas, the thymus is a single-lobed organ located cranially to the heart, in close proximity to the trachea. In many mammals (dogs, cats, non-human primates, humans), birds and fish, its location is similar, though the organ is generally bilobed (Haley, 2017). Also the histological features and composition as a lobulated organ with a distinct medulla and cortex, mainly comprised of T cells, are similar to those of most animal classes (Haley, 2003), in line with the organ’s function as a primary lymphoid organ and as the initial site for development of T cell immunological function (Abbas, 2005; Haley, 2003) In many mammals (eg. humans, non-human primates, rats, dogs) it has been shown that thymus-dependent T cell differentiation is a highly regulated process; and specific differentiation of cortical and medullary regions as well as thymic involution are biological processes that are hormonally coordinated and begin at the time of sexual maturity (Haley, 2017; Thapa and Farber, 2019). Although the function of the thymus has overall not been studied to this extent in birds, early studies in chicken have also demonstrated that the thymus produces precursors of cells involved in specific cell-mediated immune responses (T cells), and its removal impairs the ability of birds to reject allogeneic skin grafts and to produce delayed-type skin reactions (Cooper et al., 1966; Rose, 1979).

Moreover, in mammals and in birds, age-related reduction in thymic cellularity, also called thymic involution, is very apparent; here, functional tissue is replaced by adipose tissue and interlobular stroma, and epithelial structures (cords and tubules), especially of the medulla, become prominent (Ciriaco et al., 2003; Haley, 2017; Maxie, 2016; Quaglino et al., 1998). The literature regarding thymic involution in reptiles is contradicting; some authors stated that involution takes place, others claimed lifelong persistence of thymic tissue (Mader, 2009). We found no gross evidence of thymic involution in the examined boa cohort. Thymic tissue was visible also in the oldest boa, examined at 25 years of age. However, the morphometric analysis demonstrated a clear reduction in the overall cellularity of the thymus in older boas. This was due to a reduction in the size of the cortex, consistent with a loss of T cells. Older boas frequently exhibited epithelial hyperplasia and cyst formation, features found to be associated with thymic involution in mammals (Chinn et al., 2012; Pearse, 2006; Reda et al., 2019). In summary, our results indicate that some degree of thymic involution also takes place in *B. constrictor*, though this process might start late, independent of the onset of sexual hormone production (*B. constrictor* reach sexual maturity at an age of 2.5 to 3 years (Bertona and Chiaraviglio, 2003)), and progresses slowly. Moreover, our previous studies on blood haematological parameters on captive *B. constrictor* have shown that the percentage of circulating lymphocytes in the peripheral blood shows a linear association and decreases with age (Dervas et al., 2023). In diseased neonatal humans, a direct correlation between the grade of acute thymus involution and the percentage of lymphocytes in peripheral blood smears has been demonstrated (Glavina-Durdov et al., 2003). Therefore, it cannot be excluded that a possible link between thymic involution and the percentage of peripheral lymphocytes may also exist in *B. constrictor,* however this theory warrants further research.

Interestingly, we found no evidence of Hassall’s corpuscles (epithelial-derived reticular cells) in *B. constrictor.* Hassall’s corpuscles have been described in the thymus of the majority of mammals and at least in some bird and fish species, but not in amphibia (Cao et al., 2017; Kannan et al., 2015; Mikušová et al., 2017). However, cells with myofibrils and ultrastructural characteristics of striated muscle fibres were present in the thymic medulla of the boas. These cells, also called myoid cells, have also been described in many fish, amphibians, birds, and mammals, in all of which their biological role has not yet been elucidated (Varga et al., 2019).

<u>Spleen</u>. The architecture of the spleen in *B. constrictor* differed most strongly from that of other animal classes, and in particular of mammals. In boas, there was no evidence of a discernible red pulp or of periarteriolar lymphoid sheaths (PALS)/peri-ellipsoid lymphoid sheath (PELS) formation in the white pulp. This is not only in strong contrast to the spleen of mammals, birds and fish (John, 1994; Nagelkerke et al., 2018; Sayed et al., 2022) but also differs from certain other reptile species, such as the reticulated python *(Python reticulatus*), the Blotched diadem snake (*Spalerosophis diadema*) and the Schokari sand racer *(Psammophis schokari*) in which both a white and red pulp were described, although their distinction depended on the season, and the former presented as PALS/PELS-like formations (Hussein et al., 1979; Kroese et al., 1985; R. E. Ridi et al., 1981). We only examined captive snakes and can therefore not entirely rule out that the spleen has a substantially different architecture in free-living *B. constrictor* in their natural environment, however, we cannot provide a reasonable rationale for this assumption. So far, our findings indicate significant structural differences in the spleens of even phylogenetically closely related snake species, such as *B. constrictor* and pythons.

As described, the spleen of the boas lacked clear compartmentalisation and was comprised of mostly haphazardly arranged T cells, macrophage (sub)populations and a few individual B cells, IgY/IgM positive cells and plasma cells. B cells were occasionally also found in groups, vaguely reminiscent of miniscule follicles, yet without any further organisation. Instead, T cells which were mostly found disseminated throughout the organ, occasionally formed nodular (follicle-like) aggregates. These structures were not comparable with the B cell follicles known from other animal classes, with germinal centre formation; they did not even contain B cells in the centre. While B cells in birds are generated in the bursa of Fabricus, mammalian B cells appear first in embryonic liver and then, after birth, in the spleen, at the stage when haematopoiesis is taking place in these organs (Tavian and Péault, 2005). Later in the mammalian development, B cells are produced in the bone marrow and travel to the follicles (marginal zone or germinal centres) in spleen and lymph nodes, where, upon exposure to antigens, they can proliferate selectively and further differentiate to plasma cells (Oracki et al., 2010). As reptiles also lack lymph nodes and at least the *B. constrictor* seems to harbour a rather limited, structurally disorganized B cell population, the question arises as to how the reptile immune system compensates the important functions of the mammalian follicles, such as antigen presentation, B cell proliferation, plasma cell differentiation, and T cell priming (Cyster, 2010; Janeway, JR, 1989). This finding is of particular interest in light of the evolutionary theory that the increasing compartmentalization of the spleen from lower to higher vertebrates has led to separate compartments with specialized function in the immune response (Kroese and van Rooijen, 1982).

*<u>Oesophageal tonsils and GALT</u>.* While absent in mammals, oesophageal tonsils are well-known and described in birds (e.g. chicken) in which they represent part of the peripheral immune system (gut-associated lymphoepithelial tissue) (Nagy and Oláh, 2007). Histologically, avian oesophageal tonsils show a variably degree of differentiation, presenting in some bird species as a diffuse aggregate of lymphocytes while forming primary and secondary lymphoid follicles in other species; these possibly indicate different levels of maturity (Khomich et al., 2020). In the class Reptilia, oesophageal tonsils seem to represent a subtype of lymphoid tissue, possibly restricted to the family of the boid snakes (Jacobson and Collins, 1980). The evolutionary background of this phenomenon has to our knowledge not been elucidated, however, in an early study a relationship between the degree of development of lymphoid elements in the oesophagus and the dietary specialization of snakes has been suggested, with snakes that consume larger prey (rodents etc.) exhibiting more pronounced lymphoid elements (Diani A., 1974). Oesophageal tonsils were also consistently found in *B. constrictor* in the present study, where they presented as prominent, grossly evident elevated structures in the distal oesophagus that histologically consisted mostly of T cells, without any evidence of follicle formation The same was evident in the GALT, which was rich in T cells and, in contrast to many mammals (e.g. mice) and some bird species, did not show clear follicle/germinal centre formation (Jeurissen et al., 1999; Parker, 2017). Interestingly, a morphological study to characterise lymphocyte subpopulations in the oesophageal tonsils of different bird species revealed that oesophageal tonsils consist predominantly of CD20+ cells (B cells) and, at a lesser number, CD 4+ and CD8+ cells (T cells) (Khomich et al., 2020). Similarly, examination of the GALT (Peyer’s patches) in chickens demonstrated that lymphoid follicle formation occurs as early as at 7 weeks of age, with IgM^+^ and IgG^+^ cells (B cells) forming the germinal centre and CD4+ cells (T cells) located in the periphery (Kozuka et al., 2010).The immunological function of the GALT in *B. constrictor* remains to be elucidated, yet the presence of IgM expressing cells might indicate a role for the secretory immunoglobulin M in the regulation of intestinal mucosal immunity, as described in birds (Kozuka et al., 2010; Yamamoto et al., 1977).

Considering that the reptilian ancestors emerged 300 million years ago, the presumed predominance of T cells in the lymphoid tissues of the *B. constrictor* might be representative of their old phylogenetic age (Neely and Flajnik, 2016). On that note, birds, whose ancestors emerged about 225 million years ago, might represent a link between homoiothermic animals (mammals) and ectothermic animals (reptiles), which might explain their increased number of B cells and partial higher organization of the lymphoid tissue In general, the evolutionary emergence of B cell subsets and therefore also the increased diversity, specificity, and affinity of immunoglobulins is considered to be a feature of higher evolved, homeothermic vertebrates (Parra et al., 2013). This correlates with the progressive enhancement of splenic white pulp complexity and functionality over the course of vertebrate evolution (Neely and Flajnik, 2016). The relatively poor differentiation of reptile lymphoid organs compared to other animal classes might also explain why the innate immune response outweighs the adaptive response in reptiles, as for example indicated by the more limited antibody repertoire and longer time period for antibody release (Zimmerman, 2020).

### 4.2. The importance of seasonality

The majority of the studies performed on the morphology of the lymphatic tissue of reptile species (snakes, turtles) have focused on the effect of seasonal variation as they were conducted in wild animals (Hussein et al., 1979; Leceta and Zapata, 1985; R. E. Ridi et al., 1981). These studies have described distinct seasonal variations in organ size and cellularity for the thymus and the spleen, reporting a marked depletion/involution in the summer and winter, and therefore emphasizing the importance of environmental temperature when examining the immune system of ectothermic animals (Hussein et al., 1979; Leceta and Zapata, 1985; Muñoz et al., 2009; Muñoz and La Fuente, 2001; R. E. Ridi et al., 1981). Being well aware of the possibility of a direct link between the season on the haemolymphatic tissue, we took the timepoint of euthanasia into consideration when histologically evaluating the haemolymphatic tissue in the boas of the present study. The study animals were sampled at random time points throughout the year, therefore providing a representative overview and minimizing the bias. There were no histological differences in organ composition/cell types regarding the sampling season. We reasonably justified this finding by the fact that in countries like Switzerland and Germany, captive *B. constrictor* is commonly held at the same temperature throughout the year, with slight individual temperature differences between day and night and during their breeding period (as indicated in our personal communication with the breeders). In any case, the influence of seasonality and environmental factors (rainfalls etc.) - or rather the lack thereof - on the immune system of reptiles held in captivity remains a topic widely under-researched, especially in view of the reptiles’ immunocompetence/susceptibility to infectious diseases.

### 4.3. Haemolymphatic tissues in BIBD: evidence of immunosuppression?

Immunosuppression is defined as “a state of temporary or permanent dysfunction of the immune response resulting from insults to the immune system and leading to increased susceptibility to disease” (Dohms and Saif, 1984) that may or may not be associated with morphologic alterations, such as lymphoid depletion in lymphoid organs (Rahe and Murtaugh, 2017). As there is no literature on immunosuppression in reptiles, knowledge on immunosuppression in birds may be informative, as their lymphatic tissues apparently share a common past with reptiles, due to 160 million years of evolutionary dichotomy (Pope, 1991). However, a review of pathological changes of the lymphoid organs in chicken with immunosuppression due to a range of viral, bacterial (colibacillosis) and parasitic (cryptosporidiosis) infections, identified the following two reliable parameters to determine immunosuppression in poultry: 1. the number of lymphocytes in primary and secondary lymphoid organs (immunosuppression is associated with lymphocyte depletion); and 2. the number of follicular germinal centres (Pope, 1991). For instance, infection of chicken with infectious bursitis virus, a birnavirus, causes marked lymphocyte depletion in the bursa of Fabricius and the thymus. Therefore, the degree of immunosuppression in chicken can be monitored by the bursa to body weight ratio (Pope, 1991). Similarly, chicken chronically infected with protozoan parasites or infectious bursitis virus show an increase in the number of follicular germinal centres (Pope, 1991). Since, based on the results of this studies and former authors, reptiles lack classical B cell compartments, a similar approach of the “quantification” of immunosuppression in birds and reptiles is not possible. We hence tried a morphometric approach to reveal any quantitative changes in the haemolymphatic tissues in association with BIBD. This showed a significantly higher total cellularity in spleens of BIBD-positive animals vs uninfected controls; interestingly, it appeared not to be based on the increase of a specific cell type (e.g. no differences in the absolute number of macrophages/T cells), so cannot be readily explained.

The present study confirmed in snakes with BIBD that most cell types in the haemolymphatic tissues are permissive for the causative reptarenavirus, expressing viral nucleoprotein, and also bearing the typical cytoplasmic IB, without further overt cytopathic changes (Baggio et al., 2021; Hepojoki et al., 2015; Hetzel et al., 2013; Keller et al., 2017; Stenglein et al., 2012; Stenglein et al., 2017; Thiele et al., 2023; Windbichler et al., 2019), confirming the non-selective “pantropism” of reptarenaviruses. Examination of the bone marrow revealed IBs in abundant cells, including progenitor cells, at least of the erythropoietic and granulopoietic lineage. This provides evidence that reptarenaviruses infect haematopoietic stem and/or progenitor cells (HSPCs), and that the cells released into the blood stream are already infected. This is a novel finding and might explain both life-long persistence of reptarenavirus infection/BIBD and persistent/recurrent cell associated viraemia, as it is indicated by the repeated detection of IB in peripheral blood erythrocytes and leukocytes of individual snakes (personal observation). In humans, several viruses directly infect HSPCs, including cytomegalovirus, Hepatitis C virus and human herpes viruses (Pascutti et al., 2016). Such an infection appears to reduce the haematopoietic output through different mechanisms. For instance, human herpes virus 7 can infect HSPCs and impair their survival and proliferation, presumably via lysis or induced cell death (Mirandola et al., 2000). Parvovirus B19 has a selective tropism for the erythroid lineage, where productive infection induces a block in erythropoiesis that can manifest as transient or persistent erythroid aplasia (Chisaka et al., 2003). Moreover, direct infection of HSPCs by retroviruses, such as Human Immunodeficiency Virus (HIV) and Human T-cell lymphotropic virus type 1 (HTLV-1), can cause changes in the expression of intracellular factors, e.g. microRNAs, that manipulate key cellular pathways and result in the development of haematopoietic malignancies such as B cell and Hodgkin’s lymphomas (Bouzar and Willems, 2008; Neely and Flajnik, 2016).

The examination of the lymphatic tissues also revealed BIBD associated morphological changes in the spleen. Although not significantly increased in number, macrophages (Iba1+) appeared to have undergone alterations in distribution and subtype proportions with BIBD. In BIBD-positive boas, macrophages frequently formed small aggregates while they were rather diffusely distributed in control animals. In mammals, splenic macrophages are known to shape the splenic structure and/or microenvironment, and different types of macrophages and their functions have been described (Da Borges Silva et al., 2015; den Haan, Joke M M and Kraal, 2012; Nagelkerke et al., 2018). Red pulp macrophages, thought to function as a filter by facilitating clearance of senescent red blood cells and regulating iron metabolism, are most abundant and are found in the perifollicular zone (Ganz, 2012; Kurotaki et al., 2015; Mebius and Kraal, 2005). Within the Arenaviridae family, the virus most closely related to reptarenaviruses is the lymphocytic choriomeningitis virus (LCMV), a mammarenavirus causing lymphocytic choriomeningitis (and lymphocytic infiltrates in liver, adrenal, kidney, and lung), immune-complex glomerulonephritis, and vasculitis in rodents (Zhou et al., 2012). In the last decades, LCMV infection of mice has represented a useful model to dissect the mechanisms of viral persistence and generalized immunosuppression in the host (Mueller et al., 2007). Marginal zone macrophages (MZM), which toxicological studies in mice have shown to facilitate selective clearance of apoptotic cells and are able to supress immune responses to cell-associated antigens (Da Borges Silva et al., 2015; den Haan, Joke M M and Kraal, 2012; Miyake et al., 2007) appear to be an early target of LCMV; they have also been shown to be an essential cellular component in the clearance and retention of LCMV in the spleen, as they can prevent immediate virus spread to peripheral organs. Furthermore, it has been speculated that a CD8+ T cell-mediated specific destruction of LCMV-infected cells, such as macrophages and dendritic cells in secondary lymphoid organs, might lead to transient general immunosuppression in immunocompetent mice (Studstill and Hahm, 2021). This indicates that by exploiting a key element of the immune system, LCMV does induce not only a loss of antigen-presenting cells but also a disruption of the splenic microarchitecture (Müller et al., 2002). As shown, the spleen of boa constrictors lacks a clear structural organisation which precludes a functional allocation of splenic macrophages according to the microenvironment. Nonetheless, there is clear evidence of a subpopulation that stores haemosiderin and at least a mild granulomatous reaction in BIBD. We know from mammals that macrophages can proliferate locally (Amano et al., 2014); it is possible that in BIBD, certain macrophage subpopulations are replaced by others, by local loss and proliferation, respectively. This could be associated with a persistent inflammatory state in BIBD and warrants further investigations.

As mentioned, we found strong evidence of increased haemosiderosis in BIBD-positive boas, indicating increased red blood cell (RBC) resorption/destruction. This finding is of particular interest considering that mice persistently infected with certain LCMV substrains have been shown to develop an immune-mediated haemolytic anaemia that peaks around 3 weeks post infection and is associated with increased RBC clearance (Stellrecht and Vella, 1992; Stellrecht-Broomhall, 1991) and progressive loss of committed progenitor, pluripotent, morphologically differentiated cells in the bone marrow (Binder et al., 1998). The latter was attributed to excessive secretion and action of TNF-β and IFN-γ produced by CD8+ T cells (Binder et al., 1998). Interestingly, in contrast to viraemia, the LCMV-associated anaemia eventually resolved (Stellrecht and Vella, 1992). As far as we are aware from the literature and our own experience (including the present study), anaemia is not part of the disease manifestation of BIBD. However, reptarenavirus infection of pro-erythrocytes and mature peripheral RBC, together with the splenic haemosiderosis in BIBD-positive snakes indicates that the disease might indeed also be associated with alterations in erythrocyte turnover. Experimental studies are required to examine the possible effect of reptarenaviruses on the bone marrow activity.

The common co-occurrence of BIBD and secondary infections could also be confirmed in this study, as the vast majority of BIBD-positive boas showed a broad range of additional organ lesions, of which the most common was a variable degree of heterophilic (entero) colitis. The cause for the colitis was not evident upon histological examination, no infectious agents (bacteria, parasites etc.) were noted in the examined sections. Nevertheless, the heterophilic inflammation raises the question if BIBD alters the intestinal resistance to infections (also called “intestinal colonization resistance” in mammals) and therefore favours the development of opportunistic intestinal bacterial infections (Lawley and Walker, 2013) .

At last, the detailed histological examination of BIBD-positive animals detected a morphologically distinct cell type in thymus and spleen that has not been described so far. We tentatively called these “granular cells” as they contain brownish to yellowish, iron negative cytoplasmic granules that clearly differed from the granules of heterophils in HE stained sections. These cells were present in considerable numbers; they were negative for the macrophage and lymphocyte markers and were, unfortunately, not detected as part of the ultrastructural examinations of spleen and thymus. Therefore, the origin of those cells remains currently unknown. However, we have previously observed such cells in boid snakes with chronic debilitating infectious and degenerative diseases in our diagnostic case material, with or without BIBD, which would suggest that this cell type might not be a feature exclusively seen in BIBD.

## Data availability

Data will be made available on request.

## Acknowledgement

The authors wish to thank the technical staff of the Histology Laboratory and the Electron Microscopy Unit of the Institute of Veterinary Pathology for excellent technical support.

## References

Abbas, A.K., 2005. Diseases of immunity. In: Robbins and Cotran (Eds), Pathologic basis of disease, 7^th^ edition, Elsevier Saunders, Philadelphia, pp. 205–218.

Alfaro-Alarcón, A., Hetzel, U., Smura, T., Baggio, F., Morales, J.A., Kipar, A., Hepojoki, J., 2022. Boid Inclusion Body Disease is also a disease of wild boa constrictors. Microbiol. Spectr.10 (5), e0170522.

Amano, S.U., Cohen, J.L., Vangala, P., Tencerova, M., Nicoloro, S.M., Yawe, J.C., Shen, Y., Czech, M.P., Aouadi, M., 2014. Local proliferation of macrophages contributes to obesity-associated adipose tissue inflammation. Cell Metab. 19 (1), 162–171. 10.1016/j.cmet.2013.11.017.

Andreani, G., Carpenè, E., Cannavacciuolo, A., Di Girolamo, N., Ferlizza, E., Isani, G., 2014. Reference values for hematology and plasma biochemistry variables, and protein electrophoresis of healthy Hermann’s tortoises (*Testudo hermanni* ssp.). Vet. Clin. Pathol. 43 (4), 573–583. 10.1111/vcp.12203.

Argenta, F.F., Hepojoki, J., Smura, T., Szirovicza, L., Hammerschmitt, M.E., Driemeier, D., Kipar, A., Hetzel, U., 2020. Identification of reptarenaviruses, hartmaniviruses, and a novel chuvirus in captive native Brazilian boa constrictors with Boid Inclusion Body Disease. Virol. J. 94 (11). 10.1128/jvi.00001-20.

Baggio, F., Hetzel, U., Nufer, L., Kipar, A., Hepojoki, J., 2021. A subpopulation of arenavirus nucleoprotein localizes to mitochondria. Sci. Rep. 11 (1), 21048. 10.5167/uzh-209353.

Baggio, F., Hetzel, U., Prähauser, B., Dervas, E., Michalopoulou, E., Thiele, T., Kipar, A., Hepojoki, J., 2023. A Multiplex RT-PCR method for the detection of reptarenavirus infection. Viruses 15 (12), 2313. 10.3390/v15122313.

Bertona, M., Chiaraviglio, M., 2003. Reproductive biology, mating aggregations, and sexual dimorphism of the Argentine boa constrictor (*Boa constrictor occidentalis*). J. Herpetol. 37 (3), 510–516. https://www.jstor.org/stable/1566054.

Betancourt, S., Irizarry, K.J.L., Falk, B.G., Rutllant, J., Khamas, W., 2022. Micromorphological study of the upper digestive tract of the Argentine tegu (*Salvator merianae*). Anat. Histol. Embryol. 51 (2), 259–268. 10.1111/ahe.12786.

Binder, D., van den Broek, M.F., Kägi, D., Bluethmann, H., Fehr, J., Hengartner, H., Zinkernagel, R.M., 1998. Aplastic anemia rescued by exhaustion of cytokine-secreting CD8+ T cells in persistent infection with lymphocytic choriomeningitis virus. Exp. Med. 187 (11), 1903–1920. 10.1084/jem.187.11.1903.

Bortnick, A., Murre, C., 2016. Cellular and chromatin dynamics of antibody-secreting plasma cells. Dev. Biol. 5 (2), 136–149. 10.1002/wdev.213.

Bouzar, A.B., Willems, L., 2008. How HTLV-1 may subvert miRNAs for persistence and transformation. Retrovirology 5, 101. 10.1186/1742-4690-5-101.

Campbell, T.W., Grant, K.R., 2021. Exotic animal hematology and cytology, 5th edition. Wiley-Blackwell, Hoboken NJ, pp. 470–500.

Cao, J., Chen, Q., Lu, M., Hu, X., Wang, M., 2017. Histology and ultrastructure of the thymus during development in tilapia, *Oreochromis niloticus*. J. Anat. 230 (5), 720–733. 10.1111/joa.12597.

Chang, L.-W., Jacobson, E.R., 2010. Inclusion Body Disease, A worldwide infectious disease of boid snakes: A Review. J. Exot. Pet Med. 19 (3), 216–225. 10.1053/j.jepm.2010.07.014.

Cheng, H.-W., Onder, L., Novkovic, M., Soneson, C., Lütge, M., Pikor, N., Scandella, E., Robinson, M.D., Miyazaki, J.-I., Tersteegen, A., Sorg, U., Pfeffer, K., Rülicke, T., Hehlgans, T., Ludewig, B., 2019. Origin and differentiation trajectories of fibroblastic reticular cells in the splenic white pulp. Nat. Commun. 10 (1), 1739. 10.1038/s41467-019-09728-3.

Chinn, I.K., Blackburn, C.C., Manley, N.R., Sempowski, G.D., 2012. Changes in primary lymphoid organs with aging. Semin. Immunol. 24 (5), 309–320. 10.1016/j.smim.2012.04.005.

Chisaka, H., Morita, E., Yaegashi, N., Sugamura, K., 2003. Parvovirus B19 and the pathogenesis of anaemia. Rev. Med. Virol. 13 (6), 347–359. 10.1002/rmv.395.

Ciriaco, E., Píñera, P.P., Díaz-Esnal, B., Laurà, R., 2003. Age-related changes in the avian primary lymphoid organs (thymus and bursa of Fabricius). Microsc. Res. Tech .62 (6), 482–487. 10.1002/jemt.10416.

Cohen, N., 1977. Phylogenetic emergence of lymphoid tissues and cells. Marcel Dekker Inc., New York, pp. 149.

Cooper, M.D., Raymond, D.A., Peterson, R.D., South, M.A., Good, R.A., 1966. The functions of the thymus system and the bursa system in the chicken. J. Exp. Med. 123 (1), 75–102. 10.1084/jem.123.1.75.

Cyster, J.G., 2010. B cell follicles and antigen encounters of the third kind. Nat. Immunol .11 (11), 989–996. 10.1038/ni.1946.

Da Borges Silva, H., Fonseca, R., Pereira, R.M., Cassado, A.D.A., Álvarez, J.M., D’Império Lima, M.R., 2015. Splenic macrophage subsets and their function during blood-borne infections. Front. Immunol. 6, 480. 10.3389/fimmu.2015.00480.

Dabrowski, Z., Sano Martins, I.S., Tabarowski, Z., Witkowska-Pelc, E., Spadacci Morena, D.D., Spodaryk, K., Podkowa, D., 2007. Haematopoiesis in snakes (Ophidia) in early postnatal development. Cell Tissue Res. 328 (2), 291–299. 10.1007/s00441-006-0303-4.

Deldar, A., Lewis, H., Bloom, J., 1989. Electron microscopic study of the unique features and structural-morphologic relationship of canine bone marrow. Am. J. Vet. Res. 50 (1), 136–144. PMID: 2919819.

den Haan, Joke M M, Kraal, G., 2012. Innate immune functions of macrophage subpopulations in the spleen J. Innate Immun. 4 (5-6), 437–445. 10.1159/000335216.

Dervas, E., Michalopoulou, E., Liesegang, A., Novacco, M., Schwarzenberger, F., Hetzel, U., Kipar, A., 2023. Haematology, biochemistry and morphological features of peripheral blood cells in captive *Boa constrictor*. Cons. Phys. 11 (1), coad001.

Deza, F.G., Espinel, C.S., Beneitez, J.V., 2007. A novel IgA-like immunoglobulin in the reptile *Eublepharis macularius*. Dev. Comp. Immunol. 31 (6), 596–605. 10.1016/j.dci.2006.09.005.

Diani A., 1974. A comparative study of the ophidian digestive tract, Ph.D. thesis, Saint Louis University, St. Louis.

Dohms, J.E., Saif, Y.M., 1984. Criteria for evaluating immunosuppression. Avian Dis. 28 (2), 305–310. 10.2307/1590336.

Gambón-Deza, F., Espinel, C.S., 2008. IgD in the reptile leopard gecko. Mol. Immunol. 45 (12), 3470–3476. 10.1016/j.molimm.2008.02.027.

Ganz, T., 2012. Macrophages and systemic iron homeostasis. J. Innate Immun. 4 (5-6), 446–453. 10.1159/000336423.

Glavina-Durdov, M., Springer, O., Capkun, V., Saratlija-Novaković, Z., Rozić, D., Barle, M., 2003. The grade of acute thymus involution in neonates correlates with the duration of acute illness and with the percentage of lymphocytes in peripheral blood smear. J. Neonatal Biol. 83 (4), 229–234. 10.1159/000069481.

Haley, P.J., 2003. Species differences in the structure and function of the immune system. Toxicology 188 (1), 49–71. 10.1016/s0300-483x(03)00043-x.

Haley, P.J., 2017. The lymphoid system: a review of species differences. J. Toxicol. Pathol. 30 (2), 111–123. 10.1293/tox.2016-0075.

Hepojoki, J., Kipar, A., Korzyukov, Y., Bell-Sakyi, L., Vapalahti, O., Hetzel, U., 2015. Replication of boid inclusion body disease-associated arenaviruses is temperature sensitive in both boid and mammalian cells. Virol. J. 89 (2), 1119–1128. 10.5167/uzh-108061.

Hetzel, U., Sironen, T., Laurinmäki, P., Liljeroos, L., Patjas, A., Henttonen, H., Vaheri, A., Artelt, A., Kipar, A., Butcher, S.J., Vapalahti, O., Hepojoki, J., 2013. Isolation, identification, and characterization of novel arenaviruses, the etiological agents of boid inclusion body disease. Virol. J. 87 (20), 10918–10935. 10.1128/JVI.01123-13.

Holding, M.L., Frazier, J.A., Dorr, S.W., Pollock, N.B., Muelleman, P.J., Branske, A., Henningsen, S.N., Eikenaar, C., Escallón, C., Montgomery, C.E., Moore, I.T., Taylor, E.N., 2014. Wet- and dry-season steroid hormone profiles and stress reactivity of an insular dwarf snake, the Hog Island boa (*Boa constrictor imperator)*. Phys. Biochem. Zool. 87 (3), 363–373. 10.1086/675938.

Hussein, M.F., Badir, N., El Ridi, R., Akef, M., 1979. Lymphoid tissues of the snake, Spalerosophis diadema, in the different seasons. Dev. Comp. Immunol. 3, 77–88. 10.1016/s0145-305x(79)80008-7.

Imai, Y., Ibata, I., Ito, D., Ohsawa, K., Kohsaka, S., 1996. A novel gene iba1 in the major histocompatibility complex class III region encoding an EF hand protein expressed in a monocytic lineage. Biochem. Biophys. Res. Commun. 224 (3), 855–862. 10.1006/bbrc.1996.1112.

Jacobson, E.R., Collins, B.R., 1980. Tonsil-like esophageal lymphoid structures of boid snakes. Dev. Comp. Immunol. 4 (4), 703–711. 10.1016/s0145-305x(80)80071-1.

Jaffredo, T., Fellah, J.S., Dunon, D., 2006. Immunology of Birds and Reptiles. John Wiley & Sons, Ltd. 10.1038/npg.els.0000521.

Janeway, C.A., JR, 1989. The priming of helper T cells. Semin. Immunol. 1 (1), 13–20. PMID: 15630955.

Jeurissen, S.H., Wagenaar, F., Janse, E.M., 1999. Further characterization of M cells in gut-associated lymphoid tissues of the chicken. Poult. Sci. 78 (7), 965–972. 10.1093/ps/78.7.965.

John, J.L., 1994. The Avian Spleen: A Neglected Organ. Q. Rev. Biol. 69 (3), 327–351. 10.1086/418649.

Judson, J.M., Reding, D.M., Bronikowski, A.M., 2020. Immunosenescence and its influence on reproduction in a long-lived vertebrate. J. Exp. Biol. 223, pt 12. 10.1242/jeb.223057.

Kannan, T.A., Ramesh, G., Ushakumary, S., Dhinakarraj, G., Vairamuthu, S., 2015. Thymic Hassall’s corpuscles in Nandanam chicken - light and electronmicroscopic perspective (*Gallus domesticus*). J. Anim. Sci. Technol. 57 (1), 30. 10.1186/s40781-015-0064-2.

Kashimura, M., 1982. Scanning electron microscopy studies of bone marrow. Scan. Electron. Microsc.1, 445–453. PMID: 7167760.

Keller, S., Hetzel, U., Sironen, T., Korzyukov, Y., Vapalahti, O., Kipar, A., Hepojoki, J., 2017. Co-infecting reptarenaviruses can be vertically transmitted in Boa constrictor. PLoS Pathog.13 (1), e1006179. 10.1371/journal.ppat.1006179.

Khomich, V., Usenko, S., Dyshliuk, N., 2020. Morphofunctional features of the esophageal tonsil in some wild and domestic bird species. Regul. Mech. Biosyst. 11, 207–213. 10.15421/022030.

Kozuka, Y., Nasu, T., Murakami, T., Yasuda, M., 2010. Comparative studies on the secondary lymphoid tissue areas in the chicken bursa of Fabricius and calf ileal Peyer’s patch. Vet. Immunol. Immunopathol. 133 (2-4), 190–197. 10.1016/j.vetimm.2009.08.003.

Kroese, F.G., Leceta, J., Döpp, E.A., Herraez, M.P., Nieuwenhuis, P., Zapata, A., 1985. Dendritic immune complex trapping cells in the spleen of the snake, *Python reticulatus*. Dev. Comp. Immunol. 9 (4), 641–652. 10.1016/0145-305x(85)90029-1.

Kroese, F.G., van Rooijen, N., 1982. The architecture of the spleen of the Red-eared Slider, *Chrysemys scripta elegans* (Reptilia, Testudines). J. Morphol. 173 (3), 279–284. 10.1002/jmor.1051730304.

Kurotaki, D., Uede, T., Tamura, T., 2015. Functions and development of red pulp macrophages. FEMS Microbiol. Immunol. 59 (2), 55–62. 10.1111/1348-0421.12228.

Kvell, K., Cooper, E.L., Engelmann, P., Bovari, J., Nemeth, P., 2007. Blurring borders: innate immunity with adaptive features. Clin. Dev. Immunol. 2007, 83671. 10.1155/2007/83671.

Lawley, T.D., Walker, A.W., 2013. Intestinal colonization resistance. Immunology 138 (1), 1–11. 10.1111/j.1365-2567.2012.03616.x.

Leceta, J., Garrido, E., Torroba, M., Zapata, A.G., 1989. Ultrastructural changes in the thymus of the turtle *Mauremys caspica* in relation to the seasonal cycle. Cell Tissue Res. 256 (1), 213–219. 10.1007/BF00224736.

Leceta, J., Zapata, A., 1985. Seasonal changes in the thymus and spleen of the turtle, Mauremys caspica. A morphometrical, light microscopical study. Dev. Comp. Immunol. 9 (4), 653–668. 10.1016/0145-305x(85)90030-8.

Mader, R. (Ed.), 2009. Clinical anatomy of reptiles. https://www.dvm360.com/view/clinical-anatomy-reptiles-proceedings.

Maxie, M.G. (Ed.), 2016. Jubb, Kennedy & Palmer’s Pathology of Domestic Animals: Volume 3 (6th edition). W.B. Saunders, pp-102–268. 10.1016/B978-0-7020-5319-1.00013-X.

Mebius, R.E., Kraal, G., 2005. Structure and function of the spleen. Nature reviews. Immunology 5 (8), 606–616. 10.1038/nri1669.

Mikušová, R., Mešťanová, V., Polák, Š., Varga, I., 2017. What do we know about the structure of human thymic Hassall’s corpuscles? A histochemical, immunohistochemical, and electron microscopic study. Ann Anat. 2017 May;211:140–148. 10.1016/j.aanat.2017.02.006.

Mirandola, P., Secchiero, P., Pierpaoli, S., Visani, G., Zamai, L., Vitale, M., Capitani, S., Zauli, G., 2000. Infection of CD34(+) hematopoietic progenitor cells by human herpesvirus 7 (HHV-7). Blood 96 (1), 126–131. 10.1182/blood.V96.1.126 .

Miyake, Y., Asano, K., Kaise, H., Uemura, M., Nakayama, M., Tanaka, M., 2007. Critical role of macrophages in the marginal zone in the suppression of immune responses to apoptotic cell-associated antigens. J. Clin. Invest. 117 (8), 2268–2278. 10.1172/JCI31990.

Moller, C.A., Gaál, T., Mills, J.N., 2016. The hematology of captive Bobtail lizards (*Tiliqua rugosa*): blood counts, light microscopy, cytochemistry, and ultrastructure. Vet. Clin. Pathol. 45 (4), 634–647. 10.1111/vcp.12425.

Mueller, S.N., Matloubian, M., Clemens, D.M., Sharpe, A.H., Freeman, G.J., Gangappa, S., Larsen, C.P., Ahmed, R., 2007. Viral targeting of fibroblastic reticular cells contributes to immunosuppression and persistence during chronic infection. Proc Natl Acad Sci U S A. 104 (39), 15430–15435. 10.1073/pnas.0702579104.

Mukaka, M.M., 2012. Statistics corner: A guide to appropriate use of correlation coefficient in medical research. Malawi Med. J. 24 (3), 69–71. PMCID: PMC3576830.

Müller, S., Hunziker, L., Enzler, S., Bühler-Jungo, M., Di Santo, J.P., Zinkernagel, R.M., Mueller, C., 2002. Role of an intact splenic microarchitecture in early lymphocytic choriomeningitis virus production. Virol. J. 76 (5), 2375–2383. 10.1128/jvi.76.5.2375-2383.2002.

Muñoz, F.A., Estrada-Parra, S., Romero-Rojas, A., Work, T.M., Gonzalez-Ballesteros, E., Estrada-Garcia, I., 2009. Identification of CD3+ T lymphocytes in the green turtle *Chelonia mydas*. Vet. Immunol. Immunopathol. 131 (3-4), 211–217. 10.1016/j.vetimm.2009.04.015.

Muñoz, F.J., La Fuente, M. de, 2001. The effect of the seasonal cycle on the splenic leukocyte functions in the turtle *Mauremys caspica*. Physiol. Biochem. Zool: PBZ 74 (5), 660–667. 10.1086/323033.

Nagelkerke, S.Q., Bruggeman, C.W., den Haan, Joke M M, Mul, E.P.J., van den Berg, Timo K, van Bruggen, R., Kuijpers, T.W., 2018. Red pulp macrophages in the human spleen are a distinct cell population with a unique expression of Fc-γ receptors. Blood Adv. 2 (8), 941–953. 10.1182/bloodadvances.2017015008.

Nagy, N., Oláh, I., 2007. Pyloric tonsil as a novel gut-associated lymphoepithelial organ of the chicken. J. Anat.211 (3), 407–411. 10.1111/j.1469-7580.2007.00766.x.

Neely, H.R., Flajnik, M.F., 2016. Emergence and evolution of secondary lymphoid organs. Annu. Rev. Cell. Dev. Biol. 32 (1), 693–711. 10.1146/annurev-cellbio-111315-125306.

Oksanen, A., Tuurala, O., 1970. The thymus, a lymphatic and epithelial organ of the snake (*Vipera berus*). Anat. Anz. 126 (4), 400–410. PMID: 4193925.

Oracki, S.A., Walker, J.A., Hibbs, M.L., Corcoran, L.M., Tarlinton, D.M., 2010. Plasma cell development and survival. Immunol. Rev.237 (1), 140–159. 10.1111/j.1600-065X.2010.00940.x.

Parker, G.A. (Ed.), 2017. Immunopathology in toxicology and drug development, Volume 2, Organ Systems. Humana Press, Cham, Switzerland, 826 pp.

Parra, D., Takizawa, F., Sunyer, J.O., 2013. Evolution of B cell immunity. Annu. Rev. Anim. Biosci. 1, 65–97. 10.1146/annurev-animal-031412-103651.

Pascutti, M.F., Erkelens, M.N., Nolte, M.A., 2016. Impact of viral Infections on hematopoiesis: From beneficial to detrimental effects on bone marrow output. Front. Immunol. 7. 10.3389/fimmu.2016.00364.

Pearse, G., 2006. Histopathology of the thymus. Tox. Path. 34 (5), 515–547. 10.1080/01926230600978458.

Pease, D.C., 1956. An electron microscopic study of red bone marrow. Blood 11 (6), 501–526. blood.V11.6.501.501.

Pitchappan, R., Muthukkaruppan, V., 1977. Thymus-dependent lymphoid regions in the spleen of the lizard, Calotes versicolor. J. Exp. Zool. A: 199 (2), 177–188. 10.1002/jez.1401990203.

Pope, C.R., 1991. Pathology of lymphoid organs with emphasis on immunosuppression. Vet. Immunol. Immunopathol. 30 (1), 31–44. 10.1016/0165-2427(91)90006-x.

Quaglino, D., Capri, M., Bergamini, G., Euclidi, E., Zecca, L., Franceschi, C., Ronchetti, I.P., 1998. Age-dependent remodeling of rat thymus. Morphological and cytofluorimetric analysis from birth up to one year of age. Eur. J. Cell Biol. 76 (2), 156–166. 10.1016/S0171-9335(98)80029-0.

R. E. Ridi, N. Badir, S. Rouby, 1981. Effect of seasonal variations on the immune system of the snake, *Psammophis schokari*. J. Exp. Zool. 216, 357–365. 10.1002/jez.1402160303.

Rahe, M.C., Murtaugh, M.P., 2017. Mechanisms of adaptive immunity to Porcine Reproductive and Respiratory Syndrome virus. Viruses 9 (6). 10.3390/v9060148.

Raül Carmona, David M. Alba, Massimo Delfino, Josep M. Robles, Cheyenn Rotgers, Juan Vicente Bertó Mengual, Jordina Balaguer, Jordi Galindo, S. Moya Sola, 2010. Snake fossil remains from the Middle Miocene stratigraphic series of Abocador de Can Mata (els Hostalets de Pierola, Catalonia, Spain). Cidaris 30:77–84.

Reda, I., AboSalem, M.E., El Zoghby, E.M., Attia, H.F., Emam, M.A., 2019. Some histological studies on thymus gland of mature and senile rabbits. Benha Vet. Med. J. 36 (1), 218–226. 10.21608/bvmj.2019.119833.

Rose, M.E., 1979. The immune system in birds. Journal of the R. Soc. Health J. 72 (9), 701–705. 10.1177/014107687907200914.

Salakij, C., Salakij, J., Suthunmapinunta, P., Chanhome, L., 2002. Hematology, morphology and ultrastructure of blood cells and blood parasites from Puff-faced watersnakes (*Homalopsis buccata*). Kasetsart J. Soc. 36, 35–43.

Sano-Martins, I.S., Dabrowski, Z., Tabarowski, Z., Witkowska-Pelc, E., Spadacci Morena, D.D., Spodaryk, K., 2002. Haematopoiesis and a new mechanism for the release of mature blood cells from the bone marrow into the circulation in snakes (Ophidia). Cell Tis Res. 310 (1), 67–75. 10.1007/s00441-002-0557-4.

Sayed, R.K.A., Zaccone, G., Capillo, G., Albano, M., Mokhtar, D.M., 2022. Structural and functional aspects of the spleen in Molly fish *Poecilia sphenops* (Valenciennes, 1846): Synergistic interactions of stem cells, neurons, and immune cells. Biology 11 (5). 10.3390/biology11050779.

Schilliger, L., Selleri, P., Frye, F.L., 2011. Lymphoblastic lymphoma and leukemic blood profile in a red-tail boa (*Boa constrictor constrictor*) with concurrent inclusion body disease. J. Vet. Diagn. Invest, Inc 23 (1), 159–162. 10.1177/104063871102300131.

Schumacher, J., Jacobson, E.R., Homer, B.L., Gaskin, J.M., 1994. Inclusion Body Disease in boid snakes. J Zoo Wildl Med 25 (4), 511–524. 10.1128/spectrum.01705-22.

Stellrecht, K.A., Vella, A.T., 1992. Evidence for polyclonal B cell activation as the mechanism for LCMV-induced autoimmune hemolytic anemia. Immunol. Lett. 31 (3), 273–277. 10.1016/0165-2478(92)90126-9.

Stellrecht-Broomhall, K.A., 1991. Evidence for immune-mediated destruction as mechanism for LCMV-induced anemia in persistently infected mice. Viral Immunol. 4 (4), 269–280. 10.1089/vim.1991.4.269.

Stenglein, M.D., Sanchez-Migallon Guzman, D., Garcia, V.E., Layton, M.L., Hoon-Hanks, L.L., Boback, S.M., Keel, M.K., Drazenovich, T., Hawkins, M.G., DeRisi, J.L., 2017. Differential disease susceptibilities in experimentally reptarenavirus-infected boa constrictors and ball pythons. J. Virol. 91 (15). 10.1128/JVI.00451-17.

Stenglein, M.D., Sanders, C., Kistler, A.L., Ruby, J.G., Franco, J.Y., Reavill, D.R., Dunker, F., DeRisi, J.L., 2012. Identification, characterization, and in vitro culture of highly divergent arenaviruses from boa constrictors and annulated tree boas: candidate etiological agents for snake inclusion body disease. mBio 3 (4), e00180–12. 10.1128/mbio.00180-12.

Strollo, D.C., Rosado-de-Christenson, M.L., 2012. Chapter 71 - Disorders of the Mediastinum, in: Spiro, S.G., Silvestri, G.A., Agustí, A. (Eds.), Clinical Respiratory Medicine (4th edition). W.B. Saunders, Philadelphia, pp. 846–861.

Studstill, C.J., Hahm, B., 2021. Chronic LCMV infection is fortified with versatile tactics to suppress host T cell immunity and establish viral persistence. Viruses 13 (10). 10.3390/v13101951.

Tanaka, Y., Hirahara, Y., 1995. Spleen of the snake *(Elaphe climacophora*) and intrasplenic vascular architecture. J. Morphol. 226 (2), 223–235.10.1002/jmor.1052260209.

Tavian, M., Péault, B., 2005. Embryonic development of the human hematopoietic system. Int. J. Dev. Biol. 49 (2-3), 243–250.

Thapa, P., Farber, D.L., 2019. The role of the thymus in the immune response. Thorac. Surg. Clin. 29 (2), 123–131. 10.1016/j.thorsurg.2018.12.001.

Thiele, T., Baggio, F., Prähauser, B., Ruiz Subira, A., Michalopoulou, E., Kipar, A., Hetzel, U., Hepojoki, J., 2023. Reptarenavirus S segment RNA levels correlate with the presence of inclusion bodies and the number of L segments in snakes with reptarenavirus infection-lessons learned from a large breeding colony. Microbiol. Spectr. 11 (3), e0506522. 10.1128/spectrum.05065-22.

Van Haastert, Peter J M, Devreotes, P.N., 2004. Chemotaxis: signalling the way forward. Nature reviews. Mol. Cell Biol. 5 (8), 626–634. 10.1038/nrm1435.

Varga, I., Bódi, I., Kachlík, D., Mešťanová, V., Klein, M., 2019. The enigmatic thymic myoid cells – their 130 years of history, embryonic origin, function and clinical significance. Biologia 74 (5), 521–531. 10.2478/s11756-019-00214-1.

Vogel, L.A., Palackdharry, S., Zimmerman, L.M., Bowden, R.M., 2017. Humoral immune function in long-lived ectotherms, the reptiles, In: Fulop, T., Franceschi, C., Hirokawa, K., Pawelec, G. (Eds.), Handbook of Immunosenescence: Basic Understanding and Clinical Implications. Springer Cham, pp. 1–17. 10.1007/978-3-319-99375-1.

Wang, D., Tanaka-Yano, M., Meader, E., Kinney, M.A., Morris, V., Da Lummertz Rocha, E., Liu, N., Liu, T., Zhu, Q., Orkin, S.H., North, T.E., Daley, G.Q., Rowe, R.G., 2022. Developmental maturation of the hematopoietic system controlled by a Lin28b-let-7-Cbx2 axis. Cell Rep. 39 (1), 110587. 10.1016/j.celrep.2022.110587.

Windbichler, K., Michalopoulou, E., Palamides, P., Pesch, T., Jelinek, C., Vapalahti, O., Kipar, A., Hetzel, U., Hepojoki, J., 2019. Antibody response in snakes with boid inclusion body disease. PloS one 14 (9), e0221863. 10.1371/journal.pone.0221863.

Yamamoto, H., Watanabe, H., Mikami, T., 1977. Distribution of immunoglobulin and secretory component containing cells in chickens. Am. J. Vet. Res. 38 (8), 1227–1230. PMID: 562112.

Yasukazu Tanaka, Yoshihiro Hirahara, 1995. Spleen of the snake (*Elaphe climacophora*) and intrasplenic vascular architecture. Journal of morphology 226. 10.1002/jmor.1052260209.

Zapata, A., Leceta, J., Villena, A., 1981. Reptilian bone marrow. An ultrastructural study in the spanish lizard, *Lacerta hispanica*. J. Morphol. 168 (2), 137–149. 10.1002/jmor.1051680203.

Zhou, X., Ramachandran, S., Mann, M., Popkin, D.L., 2012. Role of lymphocytic choriomeningitis virus (LCMV) in understanding viral immunology: past, present and future. Viruses 4 (11), 2650–2669. 10.3390/v4112650.

Zimmerman, L.M., 2020. The reptilian perspective on vertebrate immunity: 10 years of progress. J. Exp. Biol. 223 (21). 10.1242/jeb.214171.

Zimmerman, L.M., Vogel, L.A., Bowden, R.M., 2010a. Understanding the vertebrate immune system: insights from the reptilian perspective. J. Exp. Biol. 213 (5), 661–671. 10.1242/jeb.038315.

Zimmerman, L.M., Vogel, L.A., Edwards, K.A., Bowden, R.M., 2010b. Phagocytic B cells in a reptile. Biol. Lett. 6 (2), 270–273. 10.1098/rsbl.2009.0692.

