## Supplemental files for "Haemolymphatic tissues of captive boa constrictor *(Boa constrictor):* morphological features in healthy individuals and with Boid Inclusion Body Disease"

**Supplemental Table 1.** List of animals *B. constrictor* examined as part of the study.

| Animal | Age | Sex | Haemolymphatic organs | | | | Month[2] | Further investigations[3] | Histopathological changes[4] |
|---|---|---|---|---|---|---|---|---|---|
| | | | BM | Thymus | Spleen | Alim LT | | | |
| **BIBD positive animals[1]** | | | | | | | | | |
| **A1** | 1 | M | x | x | x | x | June | Morph, BC | Focal granulomatous hepatitis, mild multifocal heterophilic enteritis |
| **A2** | 1 | M | x | x | x | x | June | Morph, BC | Mild heterophilic pneumonia and enteritis |
| **A3** | 2 | F | x | x | x | x | June | IH/ISH, morph, BC | Severe chronic diffuse epicarditis and pericarditis, mod heterophilic colitis |
| **A4** | 2 | F | x | x | x | x | June | IH/ISH, morph, BC | Mild diffuse heterophilic and granulomatous enteritis |
| **A5** | 2 | F | x | x | x | x | June | IH/ISH, morph, BC | Mild heterophilic colitis |
| **A6** | 2 | F | x | x | x | x | June | Morph, BC | Mild heterophilic colitis |
| **A7** | 2 | F | x | ne | x | x | June | Morph, BC | Mod heterophilic colitis |
| **A8** | 2 | F | x | x | x | x | June | IH/ISH, morph, BC | Mod heterophilic colitis |
| **A9** | 2 | F | x | ne | x | x | June | IH, BC | Mod heterophilic colitis |
| **A10** | 2 | M | x | ne | x | x | June | Morph, BC | Emaciation, mild heterophilic colitis |
| **A11** | 2 | M | x | x | x | x | June | ICH, morph, BC | Mild heterophilic colitis |
| **A12** | 2 | M | x | x | x | x | June | IH/ISH, morph, BC | Mild heterophilic colitis |
| **A13** | 8 | M | x | | x | x | June | IH/ISH, morph, BC | Emaciation, mod chronic active interstitial pneumonia, mild chronic interstitial nephritis, mild heterophilic enterocolitis |
| **A14** | 3 | M | x | x | x | x | June | IH/ISH, morph, BC | Mod heterophilic enterocolitis, mod heterophilic pericarditis |
| **A15** | 3 | F | x | x | x | x | June | IH/ISH, BC | Mild lymphocytic gastritis and nephritis, mild heterophilic enterocolitis |
| **A16** | 3 | F | x | x | x | x | June | BC | Mod mixed-cellular nephritis, mixed-cellular |

| | | | | | | | | | |
|---|---|---|---|---|---|---|---|---|---|
| | | | | | | | | | gastritis with fibrosis, mod mixed-cellular interstitial pneumonia |
| **A17** | 10 | F | ne | ne | x | x | June | BC | Mod to severe heterophilic colitis |
| **A18** | 10 | F | x | x | x | x | June | BC | Severe heterophilic colitis |
| **A19** | 11 | F | | x | x | ne | June | BC | Mild chronic diffuse lymphocytic tracheitis, hepatitis and interstitial pneumonia |
| **A20** | 6 | F | x | x | x | x | June | BC | Mild to mod multifocal lymphocytic interstitial pneumonia, tracheitis and nephritis, mod heterophilic colitis |
| **A21** | 11 | F | | x | x | x | June | IH, morph, BC | Mod heterophilic nephritis with mineralizations, nodular hyperplasia of the pancreas, mild chronic lymphocytic pneumonia, mild heterophilic colitis |
| **A22** | 5 | M | x | x | x | x | June | BC | - |
| **A24** | 3 | M | ne | ne | x | x | June | BC | Mild heterophilic colitis |
| **A25** | 7 | M | x | x | x | x | June | BC | Mild heterophilic colitis |
| **A26** | 4 | F | x | x | x | x | June | BC | Mild heterophilic colitis |
| **B15** | 25 | F | x | x | x | ne | August | Morph, BC | - |
| **B16** | 8 | F | ne | x | x | ne | January | IH/ISH, morph, BC | Mild chronic pneumonia and heterophilic nephritis |
| **C8** | 3.5 | F | x | ne | x | x | January | - | Mod spongiosis |
| **C9** | 1 | M | x | ne | x | x | January | - | - |
| **C10** | 1 | F | x | | | x | January | - | - |
| **C11** | 1 | F | x | x | x | x | January | IH, morph | Mild heterophilic colitis |
| **D2** | 1.5 | M | x | x | x | x | December | BC | - |
| **E** | adult | M | ne | ne | ne | ne | April | TEM | Emaciation, ulcerative dermatitis on the nose |
| **BIBD negative animals[1]** | | | | | | | | | |
| **A23** | 4 | F | x | x | x | x | June | IH/ISH, BC | Mod interstitial lymphocytic pneumonia |
| **B1** | 8 | M | x | x | x | ne | May | Morph, BC | - |
| **B2** | 8 | F | x | x | x | ne | May | BC | - |

| | | | | | | | | | |
|---|---|---|---|---|---|---|---|---|---|
| **B3** | 1 | M | x | x | x | ne | May | BC | - |
| **B4** | 10 | F | x | ne | x | ne | July | Morph, BC | - |
| **B5** | 10 | F | x | ne | x | ne | August | IH, morph, BC | - |
| **B6** | 10 | M | x | x | x | ne | August | IH, morph, BC | - |
| **B7** | 10 | M | ne | x | x | x | January | IH/ISH, morph, BC | Mild granulomatous pneumonia |
| **B8** | 6 | F | ne | x | ne | x | January | IH, moprh, BC | - |
| **B9** | 5 | M | ne | x | ne | x | January | IH, morph, BC | - |
| **B10** | 7 | M | ne | x | x | x | February | IH/ISH, morph, BC | Mod esophagitis |
| **B11** | 0.5 | M | ne | x | ne | x | February | IH, morph, BC | - |
| **B12** | 0.3 | nd | x | x | ne | x | August | TEM, BC | - |
| **B13** | adult | F | x | ne | ne | ne | September | TEM | - |
| **B14** | 0.3 | nd | x | ne | ne | ne | September | TEM, BC | - |
| **B17** | 0.3 | nd | x | x | x | x | August | TEM, BC | - |
| **B18** | 0.3 | nd | x | x | x | ne | August | TEM | - |
| **B19** | 0.3 | nd | x | ne | ne | ne | September | TEM | - |
| **C1** | 1 | nd | ne | x | x | x | September | Morph, BC | - |
| **C2** | 1 | nd | ne | ne | x | x | September | Morph, BC | - |
| **C3** | 1 | nd | ne | ne | x | x | September | IH/ISH, BC | - |
| **C4** | 1 | M | x | x | x | ne | November | IH, morph | - |
| **C5** | 1 | M | x | x | x | ne | November | - | - |
| **C6** | 1 | M | x | x | x | x | November | - | - |
| **C7** | 1 | F | x | x | x | x | November | - | - |
| **D1** | 1.5 | F | x | x | x | ne | December | IH/ISH, morph | - |

The capital letters in front of the animal numbers represent the different breeding collections or a reptile shelter . Haematopoietic tissue (bone marrow, thymus, spleen and alimentary lymphoid tissue (LT)) from the animals was subjected to morphological examinations (hematoxylin eosin stained sections (all animals). Infection with reptarenaviruses and presence of BIBD were confirmed by multiplex PCR (in all animals) and by immunohistochemistry for the viral nucleoprotein. Ne=not examined. Nd=not determined. Age: in years. Neg: negative. Pos: positive. Mod – moderate; F – female; M – male.

1 – confirmed by multiplex PCR, histological examination and immunohistochemistry for reptarenavirus nucleoprotein; 2 - month of sampling; 3 – The haemolymphatic tissues of all animals were examined histologically (HE stains and, when indicated, special stains). Further investigations were undertaken on selected animals (IH/ISH – immunohistology for leukocyte and apoptosis markers and RNA-ISH for CD20), morph - morphometric analysis; BC-buffy coat examination, TEM - transmission electron microscopy)); 4 – Histopathological changes in other organs than the haemolymphatic tissues

**Supplemental Table 2**. Morphometric analysis of BIBD -positive and negative *B. constrictor.*

<table>
<tr><th></th><th>BIBD neg<br>(Mean, n, 95%CI)</th><th>BIBD pos<br>(Mean, n, 95%CI)</th><th>Total<br>(Mean, n, 95%CI)</th></tr>
<tr><td rowspan="2">Splenic macrophages*</td><td>3.120 (8)<br>(2.086 – 4.668)</td><td>2.929 (6)<br>(0.767 – 11.188)</td><td rowspan="2">3.037 (14)<br>1.840 – 5.013)</td></tr>
<tr><td colspan="2">t=0.1296, df=12, p=0.2991</td></tr>
<tr><td rowspan="2">Splenic T cells*</td><td>27.148 (5)<br>(10.894 – 67.652)</td><td>21.00 (5)<br>(10.900 – 40.457)</td><td rowspan="2">563.77 (10)<br>(15.334 – 37.166)</td></tr>
<tr><td colspan="2">t=0.6342, df=8, p=0.5437</td></tr>
<tr><td rowspan="2">Splenic apoptotic cells</td><td>6.697 (6)<br>(2.609 – 10.785)</td><td>8.750 (4)<br>(6.652 – 10.848)</td><td rowspan="2">7.518 (10)<br>(5.241 – 9.795)</td></tr>
<tr><td colspan="2">t=-0.9992, df=8, p=0.3469</td></tr>
<tr><td rowspan="2">Splenic total cell number nuclei/area</td><td>0.022 (10)<br>(0.019 – 0.025)</td><td>0.028 (17)<br>(0.026 – 0.029)</td><td rowspan="2">0.026 (27)<br>(0.024 – 0.027)</td></tr>
<tr><td colspan="2">t=-3.7076, df=25, p<0.01</td></tr>
<tr><td rowspan="2">Thymic macrophages</td><td>2.889 (6)<br>(0.05-9.94)</td><td>4.279 (6)<br>(0.560 – 8.502)</td><td rowspan="2">3.584 (12)<br>(1.395 – 5.773)</td></tr>
<tr><td colspan="2">t=-0.6816, df=10, p=0.5110</td></tr>
<tr><td rowspan="2">Thymic T cells</td><td>44.024 (5)<br>(19.931 – 68.117)</td><td>28.392 (5)<br>(5.456 – 51.327)</td><td rowspan="2">36.208 (10)<br>(22.138 – 50.278)</td></tr>
<tr><td colspan="2">t=1.3048, df=8, p=0.2282</td></tr>
<tr><td rowspan="2">Thymic apoptotic cells</td><td>8.294 (5)<br>(3.016 – 13.573)</td><td>8.307 (5)<br>(5.904 – 10.710)</td><td rowspan="2">8.301 (10)<br>(6.073 – 10.528)</td></tr>
<tr><td colspan="2">t=-0.0061, df=8, p=0.9953</td></tr>
<tr><td rowspan="2">Thymic total cell number nuclei/area</td><td>0.024 (10)<br>(0.020 – 0.027)</td><td>0.026 (14)<br>(0.023 – 0.029)</td><td rowspan="2">0.025 (24)<br>(0.023 – 0.027)</td></tr>
<tr><td colspan="2">t=-1.2880, df=22, p=0.2111</td></tr>
</table>

*Reporting geometric means as data were subject to logarithmic transformation

Abbreviations: CI: Confidence intervals, df= Degree of freedom, neg: negative, pos: positive.

**SUPPLEMENTAL FIGURES**

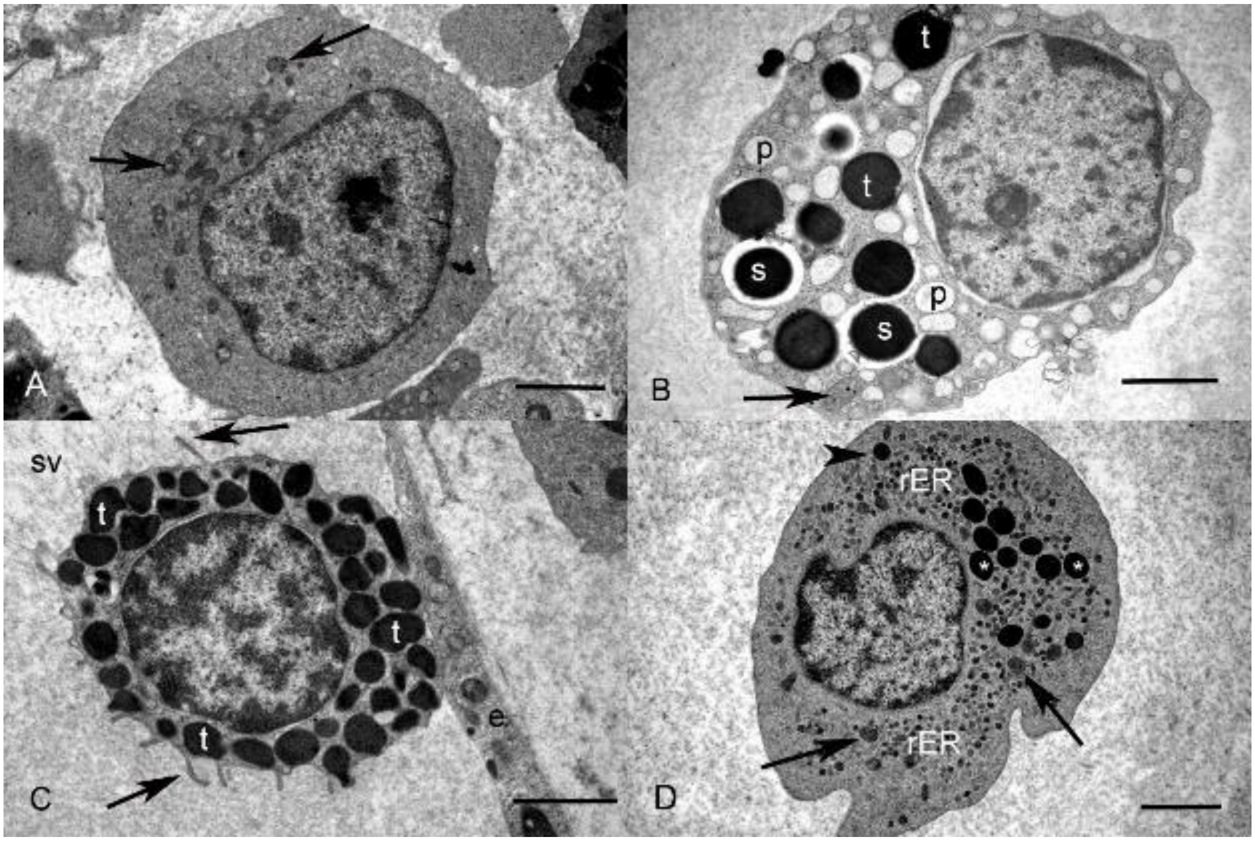


**Supplemental Fig. 1. Ultrastructural features of the haematopoietic cells in the bone marrow. A. Proerythrocyte** (animal B17, 3 months), characterised by a round central nucleus with a prominent nucleolus, a low cytoplasm:nucleus ratio, and a moderate number of mitochondria (arrows) in the cytoplasm. **B.** (**Immature) progranulocyte** (animal B18, 3 months), identified by the numerous cytoplasmic granules of very heterogenous electron density, i.e. primary granules of low electron density (p), secondary granules with electron-dense cores (s), and tertiary granules of of high electron density (t). There are a few cytoplasmic organelles (arrow: mitochondrium). The nucleus shows a prominent nucleolus and clumped, peripheral heterochromatin. **C.** (**Mature) progranulocyte** (animal B18, 3 months) in the lumen of the sinus venosus (sv), attached to an endothelial cell (e). It contains abundant electron dense tertiary granules (t) and has cytoplasmic projections (pseudopodia, arrows). **D. Promonocyte** (animal B18, 3 months)**,** characterised by s slightly eccentric and slightly indented nucleus with a moderate amount of peripheralized heterochromatin. The cytoplasm is abundant and contains a high number of organelles, i.e.

mitochondria (arrow), rough endoplasmic reticulum (rER) and lysosomes (arrowhead), as well as a moderate number of small electron-dense granules (asterisks). TEM, bars = 2 µm.

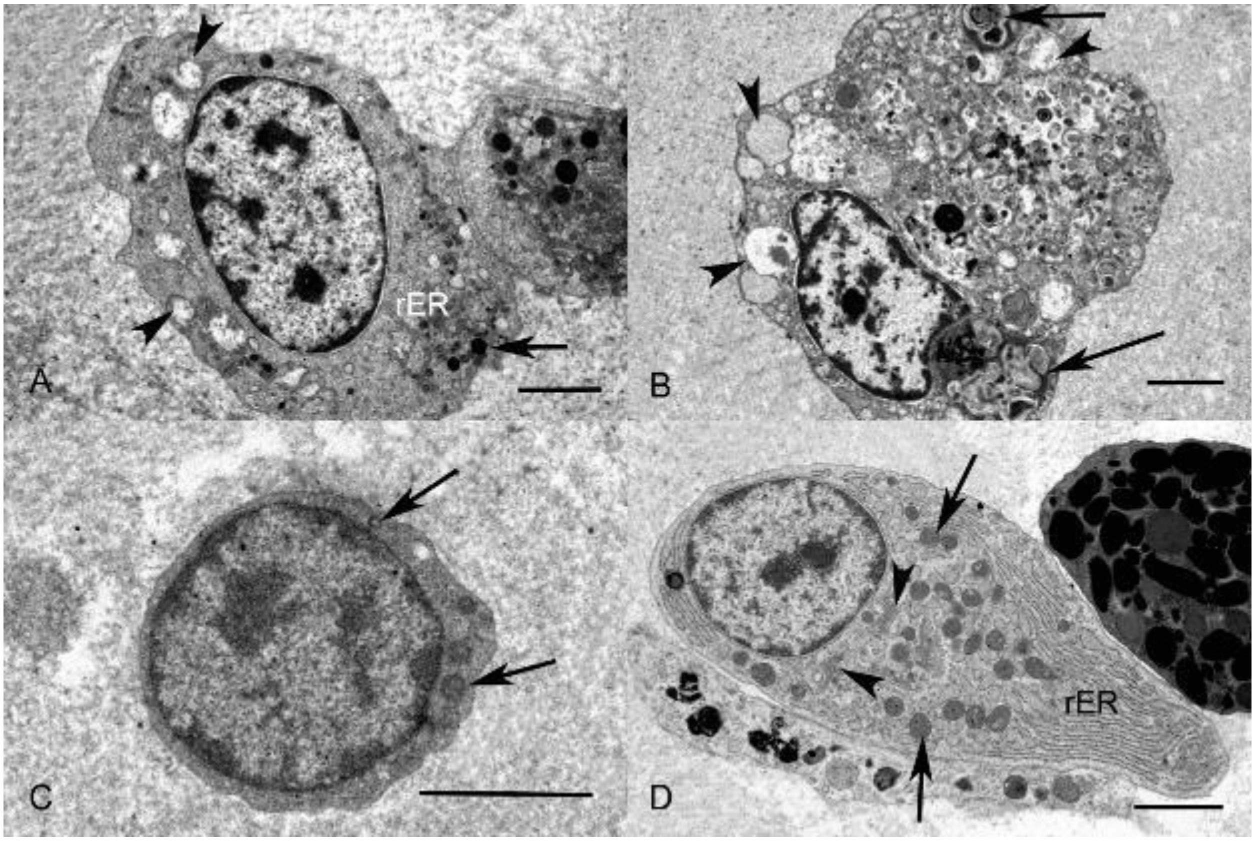


**Supplemental Fig. 2. Ultrastructural features of bone marrow cells. A-C. Precursor cells** (animal B17, 3 months). **A. Proazurophil,** characterized by a cytoplasm that contains variably sized vacuoles filled with finely granular material of low electron density (arrowheads), and a few organelles (rough endoplasmic reticulum and lysosomes (arrow)). The nucleus is roundish, with a moderate amount of clumped heterochromatin. **B. Proazurophil.** The cytoplasm is filled with abundant vacuoles that contain finely granular material of low electron density (arrowheads) and irregularly shaped, lamellar, myelin-like material (arrows). **C. Prolymphocyte**. The cell contains a large, central round nucleus with clumped heterochromatin and a small rim of cytoplasm (arrows: mitochondria). **D. Plasma cell** (animal B18, 3 months)**.** The cell is oval, with an eccentric round nucleus. The heterochromatin forms central clumps and is radially arranged in the periphery ("clockface" appearance). The cytoplasm is filled with rough endoplasmic reticulum (rER) organized in closely spaced cisternae, several mitochondria (arrows) and a Golgi apparatus (arrowheads). Bars = 2 µm.
